# The conserved landscape of RNA modifications and transcript diversity across mammalian evolution

**DOI:** 10.1101/2024.11.24.624934

**Authors:** G Santos-Rodriguez, A Srivastava, A Ravindran, F Oyelami, KH Ip, P Gupta, J Villanueva, HE King, A Grootveld, J Blackburn, I Gupta, HGS Vieira, NE Shirokikh, E Eyras, RJ Weatheritt

## Abstract

Gene expression programs underpin the development of shared phenotypes, yet the importance of transcript complexity in shaping mammalian evolution remains unclear. Here we present a comprehensive long-read direct RNA sequencing atlas of full-length transcripts and their m6A modifications across six tissues (hippocampus, frontal cortex, cerebellum, testes, skeletal muscle, and liver) in five mammals (human, mouse, rat, dog, and cow) and a non-mammalian out-group (chicken). Our analysis reveals that 29% of genes have multiple mammalian-conserved alternative transcripts, with 31% of these genes showing tissue-specific switching of the major transcript isoforms. We uncover extensive conservation of coordinated splicing events, primarily driven by tissue-specific mutually associated exon splicing, particularly in neural tissues and cytoskeletal genes. At the epitranscriptome level, we find that 14.2% of m6A RNA modifications are conserved across mammals, with 39% of analysed genes containing a conserved m6A RNA modification. Our work provides unprecedented insight into the evolution of transcript complexity and the epitranscriptome, highlighting their potential roles in shaping mammalian phenotypic diversity and providing a valuable resource for understanding post-transcriptional regulation across mammalian evolutionary time.

## Introduction

Transcriptome complexity in mammals is driven by two key mechanisms: the dynamic regulation of transcript composition and the post-transcriptional deposition of RNA modification^1-3^. Together, these processes generate an immense diversity of isoforms across mammalian species, contributing to phenotypic complexity^4,5^. However, the extent of transcript complexity conservation and its role in shaping mammalian evolution remain unclear.

Transcript sequence composition is primarily regulated through alternative splicing, alternative promoter usage, and alternative polyadenylation^1,6,7^. These processes are highly controlled and pervasive in mammals, impacting most multi-exonic genes^2,8^. While multiple studies have demonstrated that the expansion of transcript complexity drives mammalian-specific phenotypes^4,5^, genome-wide analyses have suggested that only a small proportion of individual alternative splicing events show evolutionary conservation across mammals^9,10^. However, these studies were restricted by the limitations of short-read sequencing, focusing primarily on individual exons rather than full-length transcripts. Consequently, the extent of conservation for complete mRNA transcripts and the coordination of alternative splicing, promoter, and polyadenylation events within genes across mammalian evolution remains unknown.

Transcript complexity is further enhanced by the dynamic deposition of RNA modifications onto mRNA molecules, forming the epitranscriptome^3^. Over 150 RNA modifications have been identified, with N6-methyladenosine (m6A) being the most prevalent in human mRNAs^11,12^. These modifications contribute to gene regulation by impacting mRNA stability, localization, and translation efficiency^13,14^. However, our understanding of the roles of RNA modifications in mammalian evolution has been limited by the lack of genome-wide approaches capable of determining their conservation at nucleotide resolution across multiple species.

Long-read direct RNA sequencing technologies now enable the analysis of full-length transcripts while preserving the modifications present in RNA^15,16^. This technological advancement provides an unprecedented opportunity to investigate the conservation of transcript complexity and RNA modifications across mammalian evolution at high resolution.

Here, we present a comprehensive atlas of full-length transcripts and their m6A RNA modifications for six tissues (hippocampus, frontal cortex, cerebellum, testes, skeletal muscle, and liver) across five mammalian (human, mouse, rat, dog, and cow) and a non-mammalian species (chicken). Using long-read direct RNA sequencing, we uncover extensive conservation of alternative transcripts and coordinated splicing events across mammals. We further reveal the evolutionary dynamics of m6A modifications, identifying a subset of highly conserved sites with distinct characteristics. Our findings provide new insights into the evolution of transcript complexity and the epitranscriptome, highlighting their potential roles in shaping mammalian phenotypic diversity.

### Evolution of transcript complexity

Long-read direct RNA nanopore sequencing data were collected from cerebellum, hippocampus, frontal cortex, liver, skeletal muscle, and testis across five mammalian species (human, mouse, rat, dog, cow) and a non-mammalian out-group (chicken) (Figure 1A and S1A-D; Table S1; Methods) covering approximately 320 million years of evolution. To compensate for differences in annotation quality between species and to ensure a robust comparison, for each species, all annotated transcripts from the Ensembl database^17^ were considered and supplemented with orthologous human transcripts based on the conservation of orthologous exon structure (Methods and Figure S1E; Tables S2, S3). These extended transcriptomes establish orthologous relationships between genes and transcripts, enabling direct species comparisons. Reads were mapped to the extended transcriptomes to quantify gene and transcript expression, with the high mapping percentage validating the approach of creating extended transcriptomes (Figure S1B and Table S1).

**Figure 1.**
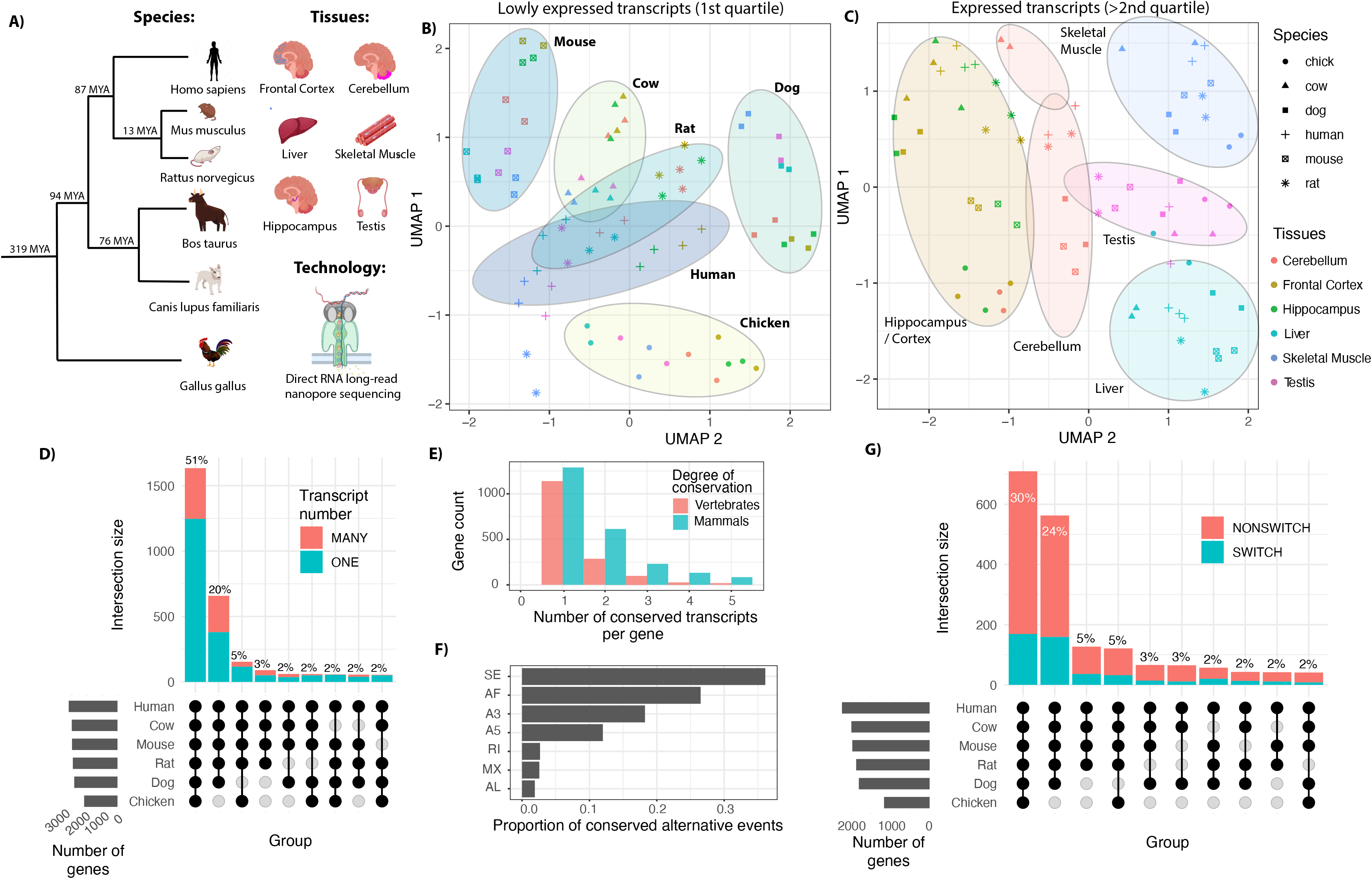
**(A)** Overview schematic of experiment with phylogenetic tree detailing evolutionary distance in millions of years (MYA) between species surveyed on left and tissues on right**(B)** UMAP plot of transcripts in lower quartile of relative expression (< 25%). Datasets are indicated by species-specific labels. **(C)** UMAP plot of transcripts with high relative expression (>25%). Datasets are indicated by tissue-specific labels. **(D)** A cumulative UpSet plot describing the conservation of genes with a single or multiple isoforms. An UpSet plots the intersections of a set as a matrix. Each column corresponds to a set, and bar charts on top show the size of the set. Each row corresponds to a possible intersection: the filled-in cells show which set is part of an intersection. **(E)** A histogram describing the number of transcripts per gene conserved across mammalian and vertebrate species, respectively. **(F)** Proportion of alternative splicing events leading to differences between conserved mammalian transcripts. SE = skipping exon; AF = alternative first exon; A3 = Alternative 3’ splice site, A5 = Alternative 5’ splice site; RI = retained intron; MX = mutually exclusive exons; AL = alternative last exon. **(G)** A cumulative UpSet plot describing extent of conserved switching between tissues. A switch is defined as a tissue-specific change if it occurs in the transcript with highest relative expression in that tissue.

To provide an overview of the pan-vertebrate long-read dataset, non-expressed genes and low abundance transcripts (CPM < 1) were removed (Methods; Table S4) The remaining transcripts were then binned based on their abundance relative to gene expression (percentage of transcript abundance per gene, PTE). Finally, transcript expression was visualized using dimensionality reduction analysis. Notably, transcripts in the lowest quartile of expression cluster in a species-specific manner (Figure 1B). In contrast, transcripts from all other quartiles of total gene expression display tissue-specific clustering patterns (Figure 1C). To quantify species-specific expression, the fraction of detected transcripts (CPM > 0) unique to the human transcriptome was assessed. Consistent with previous evolutionary studies showing limited conservation of alternative splicing at the exon level^9,18^, 71.0% (25,295/35,585) of transcripts in the dataset are species-specific. A similar fraction of species-specific transcript expression is observed across all investigated species (Figure S1F). Despite their large numbers, these species-specific transcripts are of low abundance, with each transcript contributing a median of just 4.26% to its total gene expression when expressed (Figure S1G). This observation aligns with previous observations based on individual exons^19^, where species-specific exons often have low and noisy expression.

The expression conservation of transcripts with conserved exon-intron structure (termed here as “structurally conserved transcripts”) across vertebrate evolution was next investigated. The analysis revealed that mammalian conserved transcripts comprise 17.7% (4,044/22,791) of the structurally conserved transcripts. In contrast to the relatively small contribution of species-specific transcripts, each structurally conserved transcript contributes a median of 28.7% percentage to total gene expression (Figure S1G). This translates into 71.0% (2,372/3,341) of the genes analysed having one or more transcripts conserved across mammals (Figure 1D). Furthermore, 29.2% (978/3,341) of genes have multiple conserved transcripts, with 11.3% (378/3,341) having 3 or more transcripts conserved across mammals (Figure 1D). Expanding our analysis to non-mammalian species, 13.5% (428/3,173) of genes have multiple conserved transcripts, with 3.97% (126/3,173) having three or more conserved transcripts (Figure 1E). Examination of the drives of the differences in transcript composition between these conserved transcripts revealed that changes are primarily due to differential regulation of internal exons by alternative splicing and first exons by alternative promoter usage (Figure 1F).

Given that tissue-specific alternative splicing events are major drivers of phenotypic diversity^6^], the frequency of genes with multiple transcripts exhibiting tissue-specific switches in major transcript isoforms was investigated. We observe that for genes with multiple conserved transcripts, 37.4% (160/428) in vertebrates and 31.1% (304/978) in mammals display tissue-dependent switching of the major transcript (Figure 1G). Notably, these switch-like events are especially enriched in cytoskeletal and synaptic proteins (Figure S1I), underscoring the importance of alternative transcripts in driving cellular diversity in these processes across mammalian evolution.

### Conservation of coordinated splicing

A major advantage of the long-read dataset is that it produces reads covering multiple exons, often spanning entire transcripts (Figure S1C-D). The nature of this data enabled us to investigate the coordination of alternative splicing, alternative promoter usage, and alternative polyadenylation events, all of which are essential for generating mature mRNA transcripts^20-22^. This contrasts with the limited exploration of these intricate relationships using short-read sequencing techniques, where read length rarely spans more than two exons^20-22^. Consequently, no prior study has examined the conservation of alternative exon coordination across the evolutionary spectrum of mammals and vertebrates.

To accurately detect and quantify all coordinated events, all possible combinations of exons within transcripts supported by more than 10 reads were considered. Among the associated exon pairs, those whose co-regulation had a false-discovery rate of less than 0.05 were considered coordinated (Methods). These pairs were categorised into two distinct groups: adjacent exons (consecutively coordinated exons) and distant exons (coordinated pairs separated by at least one constitutive exon) (Figure 2A and Table S5). Initial analysis of transcripts shows that 25.4% (666/2,622) of assessed genes with multiple expressed transcripts have conserved coordinated splicing events, with 58.0% (5,099/8,785) of these exons involved in adjacent splicing events. These coordinated events are primarily associations between internal exons (Figure 2B). However, conserved distant events are enriched for start-end exons (Figure 2C, Chi-squared p < 4.29 × 10^−04^, odds ratio = 1.52, as compared to non-conserved events), whereas conserved adjacent events are enriched for internal exons (Figure 2C, p < 2.72 × 10^−06^, Chi-squared test). Collectively, these results demonstrate that coordinated splicing is highly conserved across mammalian species.

**Figure 2.**
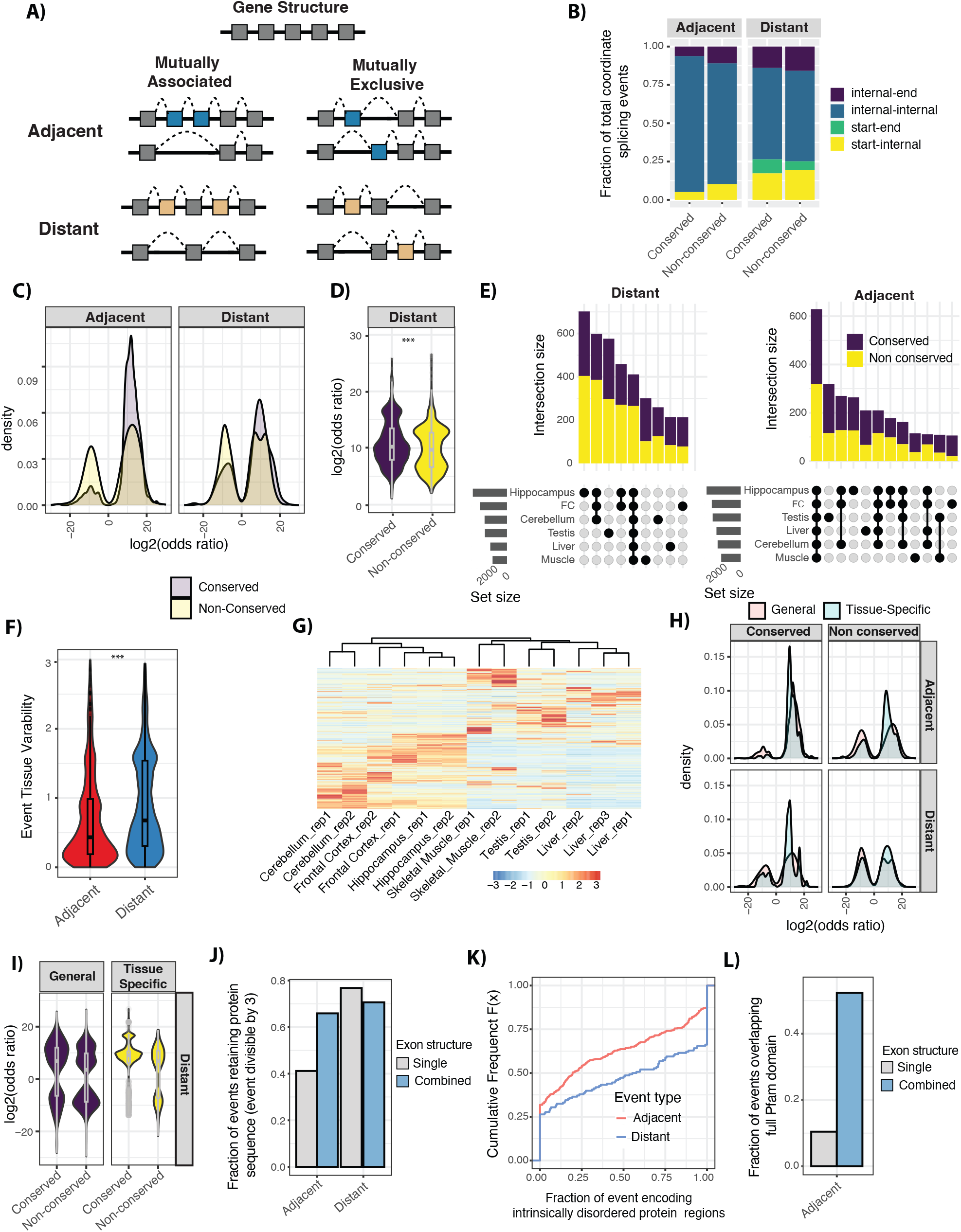
**(A)** A schematic defining the distinct types of coordinated splicing events investigated. Alternative exons involved in adjacent splicing events are shown in blue. Alternative exons involved in distant splicing events are shown in yellow. Exons that are constitutive are shown in grey. **(B)** A cumulative bar plot of relative position within a transcript of adjacent (left) and distant (right) coordinated splicing events within conserved and non-conserved transcripts. **(C)** Density plots for the log-odds ratio for adjacent and distant exon pairs. Positive and negative logs-odds ratio represent mutually associated and mutually exclusive coordinated events, respectively. **(D)** A violin plot describing the log-odds ratio for conserved and non-conserved mutually associated coordinated splicing events. A violin plot depicts distributions of numeric data for one or more groups using density curves. The width of each curve corresponds with the approximate frequency of data points in each region. (**E**) A cumulative UpSet plot describing the tissue-specificity of distant (left) and adjacent (right) coordinated splicing events with shading of bar plots illustrating the conservation of transcripts. Refer to Figure 1D legend for an explanation of an UpSet plot. (**F**) A violin plot describing the tissue-specific exon inclusion variability of alternative exons involved in adjacent and distant coordinated splicing events. Refer to Figure 2D legend for an explanation of violin plots here. **(G)** A heatmap of normalised expression for transcripts with conserved distant coordinated splicing events across all tissues sequenced. **(H)** Density plots for the log-odds ratio for tissue-specific and non-tissue-specific adjacent and distant coordinated exon pairs. Positive and negative logs-odds ratio represent mutually associated and mutually exclusive coordinated events, respectively. **(I)** Violin plots describing the log-odds ratio for tissue-specific and non-tissue-specific conserved and non-conserved distant coordinated splicing events. **(J)** A bar plot of the fraction of events that maintain the open reading frame when included in transcript when exons are considered individually (single) or as part of a complete coordinated event (combined); (n=973, 1,099). **(K)** A Cumulative distribution plot of the fraction of exons encoding intrinsically disordered regions for adjacent and distant coordinated splicing events. **(L)** A bar plot of the fraction of events encompassing a full Pfam domain when exons are considered individually (single) or as part of a complete adjacent coordinated event (combined).

The next step was to investigate whether exons involved in conserved coordinated splicing events tended to co-occur in transcripts (mutually associated) or appear exclusively of one another (mutually exclusive), as shown in Figure 2A. Consistent with previous studies^22^, distant exon pairs demonstrate an increased tendency for mutually associated coordination (Figure 2C). This trend is accentuated when events are required to be conserved across mammalian species, with conserved distant events displaying an even more pronounced enrichment for mutually associated distant events (Figure 2B, p < 6.50×10^−50^, Wilcoxon rank-sum test), as well an increased strength of coordination (Figure 2D, p < 8.59 × 10^−12;^ Figure S2C, p < 2.41 × 10^−12^, Wilcoxon rank-sum test). A similar trend was observed for adjacent events (Figure 2B, p < 2.27×10^−48^, Wilcoxon rank-sum test).

The tissue expression patterns of conserved coordinated splicing events were next investigated. Initially, the expression profiles of both conserved and non-conserved coordinated splicing events across analysed tissues were considered. This analysis revealed a striking divergence in the tissue specificity between distant and adjacent coordinated splicing (Figure 2E). Specifically, distant events show a tendency for tissue-specific expression, while adjacent events display a more tissue-wide expression pattern (Figure 2E). To quantify this divergence, tissue-specific gene expression variability based on the percentage of transcript expression per gene (Methods) was calculated. This analysis shows that distant splicing events are significantly enriched for highly variable events, underscoring their pronounced tissue-specificity (Figure 2F, p < 7.47 × 10^−36^, Wilcoxon rank-sum test). Furthermore, wherever the conservation status of these events is identified, a substantial enrichment of conserved splicing events is observed among tissue-specific events (Figure 2E, p < 6.50×10^−50^, Wilcoxon rank-sum test).

This tissue-variability of distantly coordinated splicing is most pronounced in the comparison between neural and non-neural tissue types (Figure 2G) and is driven by genes with specialised neuronal functions, such as those involved in actin processes and cytoskeletal organisation (Figure S2E). Further investigation of tissue-specific events reveals a strong enrichment for mutually associated splicing (Figure 2H, p < 5.76×10^−32^, Fisher-exact test). This enrichment is significantly more pronounced than that observed in conserved non-tissue-specific distant events (Figure 2I, p < 8.59×10^−12^, Fisher-exact test). Together, these results demonstrate the prevalence and importance of tissue-specific mutually associated coordinated splicing in mammalian evolution.

Finally, the potential impacts of coordinated splicing events on the proteome were examined by observing alternative splicing events transcript variation within the coding sequences. Consistent with their enrichment in tissue-specific exons^23^, distant coordinated events exhibit a strong tendency to preserve the open reading frame and encode intrinsically disordered regions. In contrast, adjacent exons show no evidence of a selection to maintain the open reading frame (Figure 2J). However, when exons within adjacent coordinated splicing events are considered together, there is a strong tendency to maintain the open reading frame (Figure 2J, p < 5.11 × 10^−10^, Fisher exact test), underscoring the importance of considering the coordinated regulation of exons. Since adjacent coordinated events collectively tend to maintain the open reading frame, we next evaluated how the contributing exons affect protein features. Notably, these events are enriched in structured regions of the respective proteins (Figure 2K, p < 8.32 × 10^−07^, Fisher exact test; Methods). However, only 10% of exons (836 / 8,012) involved in adjacent splicing events overlapped a full Pfam structured domain suggesting the majority would disrupt protein folding. When considered as adjacent coordinated events were considered all together, over 50% of these events (1,654 / 3,159) lead to the complete removal of a structured domain from the respective protein (Figure 2L, p < 2.18 × 10^−75,^ Fisher exact test). These findings demonstrate that coordinated splicing has, to a significant degree, evolved to uphold the structural integrity of the encoded proteins. This underscores the importance of considering coordinated splicing events when assessing the impact of alternative splicing on proteomic complexity.

### Conservation of m6A epitranscriptome

Nanopore direct RNA sequencing enables the analysis of native RNA molecules without the need for reverse transcription. This approach preserves epitranscriptomic modifications present in the RNA, allowing for their detection and characterisation. To accurately detect and quantify all m6A modification sites in our dataset across species, we used an artificial intelligence (AI)-based method ^15^, capable of detecting RNA modifications and their stoichiometry at single-nucleotide resolution. To account for inter-species detection rate differences, we included only adenosines observed with >20 read coverage in at least one tissue across all species (Figures S3-7; Table S6; Methods). Initial clustering of all m6A sites with conserved adenosine tested in all species reveals that m6A modifications tend to be species-specific (Figure 3A). However, evidence of conservation is demonstrated by the phylogeny-based clustering of species, with the out-group chicken dataset as an outlier (Figure 3A). Analysis of the localisation of m6A modification sites within transcripts confirms previous observations^12,24,25,12,24,25^ that m6A modifications are relatively enriched near stop codons and depleted at exon-exon junctions (Figures 3B-C).

**Figure 3.**
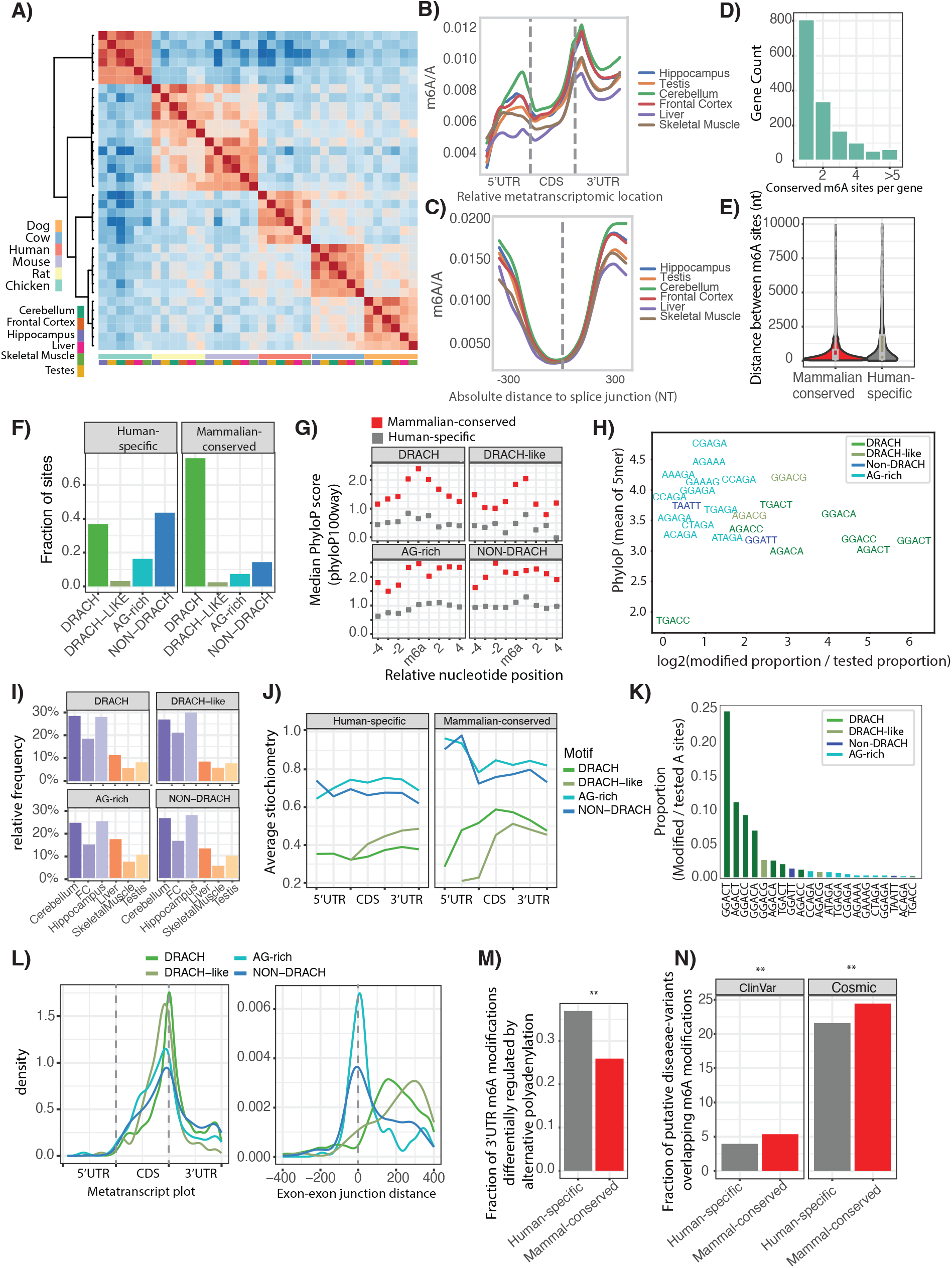
**(A)** A correlative heatmap of the conservation of all detected m6A modifications across sites tested in at least one tissue in every species. **(B)** A metatranscript profile of the relative density of m6A modifications across mRNA regions. **(C)** A metatranscript relative density profile of the m6A modifications surrounding exon-exon splice junctions. **(D)** A histogram displaying the number of conserved m6A sites per gene. **(E)** A violin plot describing the absolute genomic distance between conserved (red) and human-specific (grey) m6A modifications in a gene body. Refer to Figure 2D legend for an explanation of violin plots here. **(F)** A bar plot depicting the fraction of m6A sites assigned to motif classes for human-specific m6A sites (left panel) and conserved m6A sites (right panel). **(G)** A dot plot describing the conservation of nucleotides (PhyloP score) surrounding conserved and human-specific m6A modifications in 3’UTR. Only sites with the central adenosine conserved and tested across all species were included. **(H)** A scatter plot describing the PhyloP conservation (y-axis) and the relative usage (x-axis) of 5-mer motifs surrounding conserved m6A sites. The motifs with >=5 m6A sites were considered. **(I)** A bar plot describing the relative frequency of m6A usage for each motif class across tissues. **(J)** A metatranscript density profile plot of the average m6A stoichiometry based on transcript position of human-specific (left) and conserved (right) m6A sites, separated by motif type. **(K)** A bar plot showing the proportion of tested sites displaying a conserved m6A site (y-axis) separated by 5-mer motif (x-axis) (DRACH, DRACH-like, AG-rich motif, and rest of non-DRACH motifs). The motifs with >=5 m6A sites were considered. The average of these proportions for DRACH motifs is 5.5%. Bar plot is coloured based on the motif match. **(L)** A metatranscript density plot of the position of conserved m6A sites across transcripts (left panel) and exon-exon junctions (right panel), separated by motif type. **(M)** A bar plot for the fraction of human-specific and conserved m6A sites that occur in isoform-specific regions of the 3’UTR. **(N)** Bar plots of the fraction of human-specific and conserved m6A sites that overlap non-benign germline variants from ClinVar and somatic variants from COSMIC.

We next examined the conservation of m6A sites, defining conserved modifications as those present in human and at least two other species while within the same tissue type (Figures S8-10; Table S7). Despite the species-specific clustering of m6A sites^26^, our analysis reveals that 14.2% of m6A RNA modifications are conserved with this percentage increasing to 20.5 % when only m6A modifications overlapping DRACH motifs were considered. Moreover, 38.6% of analysed genes (1,909 out of 4,945) contain at least one conserved RNA modification. In fact, 18.1% (897 out of 4,945) of analysed genes have multiple conserved m6A modifications (Figure 3D). Comparing the positioning of conserved sites with human-specific m6A modifications (Tables S8) reveals that conserved sites cluster more closely together. The median distance between conserved sites is just 342 nucleotides in genomic space, with this clustering primarily occurring in the 3’-UTR (Figure 3E, p < 5.75 × 10^−16^, Wilcoxon rank-sum test).

Previous studies have proposed that m6A deposition is primarily driven by the presence of the DRACH sequence motif^24^. Consistent with this, the proportion of m6A sites matching the DRACH motif increases with the number of species where conserved m6A sites are detected (Figure S11A), with 78.3% of conserved m6A sites matching the DRACH motif (Figure 3F and Table S8). While DRACH motifs are the most common, notably, 21.7% of conserved m6A sites do not match the DRACH sequence (Figure 3F and S11B). Further analysis of these conserved m6A sites reveals enrichment for AG-rich motifs, as well as the presence of DRACH-like (e.g., AGAGC) and non-DRACH (e.g., GGATT) motifs (Figure 3F and Table S8).

To orthogonally validate the conservation of DRACH, DRACH-like, AG-rich and non-DRACH m6A sites, we assessed the conservation of nucleotides surrounding the conserved m6A sites compared to human-specific sites. This analysis demonstrates significantly higher conservation (PhyloP score) in the 9-mer context for all motif types compared to human-specific sites (Figure 3G, DRACH: p < 3.13 × 10^−180^; DRACH-like: p < 2.33 × 10^−10^; AG-rich: 9.00 × 10^−44^; non-DRACH: p < 2.44 × 10^−126^; Wilcox rank-sum test). Interestingly, the 9mer context of the conserved AG-rich and non-DRACH m6A is more strongly conserved than DRACH motifs (Figure 3G) (AG-rich: p < 7.56 × 10^−09^; non-DRACH: p < 7.45 × 10^−25^, Wilcox rank-sum test). This observation remains consistent regardless of the specific DRACH motif variant, tissue type, or modification frequency (Figures 3H, 3I and S11D).

The relative occurrence of m6A sites within transcripts is quantified using stoichiometry, where higher values indicate more frequent usage of a site. Conserved m6A sites consistently display increased stoichiometry levels compared to human-specific sites (Figure 3J, Figure S11C, DRACH: p < 2.68 × 10^−109^; DRACH-like: p < 0.076; AG-rich: p < 0.019; non-DRACH: p < 2.78 × 10^−33^; Wilcox-rank sum). More surprisingly, both non-DRACH and AG-rich m6A sites display significantly higher stoichiometry levels than m6A modifications at DRACH motifs (Figure 3J, AG-rich: p < 5.77 × 10^−12^; non-DRACH: p < 9.80 × 10^−148^). This is despite DRACH motifs being more frequently modified (Figure 3K). Further evaluation of the modification frequency of conserved adenosine within DRACH motifs shows that only 5.5% of these sites are consistently modified across species and tissues (Figure 3K). This percentage increases only slightly when regions around exon-exon junctions are excluded (Figure S11E).

To evaluate potential differences between these two m6A types, we analysed their relative occurrence within transcripts. Conserved m6A types are enriched at the stop codon compared to human-specific sites (Figure 3L, left panel, Figure S11F, DRACH: p < 4.73 × 10^−04^; non-DRACH: 4.74 × 10^−05^; Fisher-exact test). On the other hand, while m6A DRACH and DRACH-like sites are depleted at exon-exon junctions (p < 1.35 × 10^−07^, Fisher exact test), non-DRACH and AG-rich sites show a relative enrichment around these positions (Figure 3L, right panel, Figure S11G-H, non-DRACH: p < 1.41 × 10^−09^; AG-rich: p < 1.10 × 10^−06^, Fisher-exact test).

A key advantage of detecting m6A sites from nanopore data is the ability to evaluate the relationship of these modifications with transcript composition and expression. Given the enrichment of m6A sites in the 3’UTR, we evaluated the isoform-specificity of these modifications. We focused on variation due to alternative polyadenylation site selection and alternative last exon usage of sufficiently well covered transcripts (coverage > 5). This analysis reveals that, on average, 32.0% (611 / 1,910) of m6A sites within 3’UTR display transcript-specific expression. These transcript-specific m6A sites are enriched for human-specific modifications (Figure 3M, p < 0.0010, Fisher exact test). Nevertheless, 27.3% (240 of 878) of conserved modifications also display transcript-specific regulation, suggesting this type of regulation is widespread. Finally, we assessed the occurrence of m6A sites over putative disease-related mutations using data from ClinVar and COSMIC. Consistent with the biological significance of m6A modifications, conserved m6A sites more frequently overlap with disease-related mutations (Figure 3N, p < 0.0021, Fisher exact test). This suggests that disruption of m6A deposition may play a role in disease pathogenesis.

## Discussion

Our comparative analyses of transcript complexity across 6 species reveals a stronger transcript-level conservation across evolution than previously reported for individual exons^9,10^. This discrepancy is likely because the majority of alternatively spliced exons are included at low levels^9,18,19^, dominating the incidence of alternative splicing across evolution. The prevalence of this unproductive splicing^27^ is likely under-estimated in this paper, as our focus on conservation biased towards annotated exons and conserved exon-exon structures. Thus, future analysis of our dataset may help also reveal extensive species- and lineage-specific transcript patterns that may drive phenotypic complexity^4,5^. In terms of conserved transcript structures, our analysis and use of long-read nanopore sequencing reveals that over two-fifths of human genes have alternative transcripts conserved across mammalian evolution, with many demonstrating tissue-specific inclusion in adult tissues. Given the extensive evidence of dynamic alternative splicing switches in early development^28,29^, this number is likely to be a minimum baseline for the conservation of alternative transcripts.

Our work also provides insight into the conservation of coordinated splicing across evolution, revealing it across both local and distant splicing events. For distant coordinated splicing events, we demonstrate tissue-specific regulation, especially for neural tissues and in cytoskeletal genes. This finding aligns with previous studies from short-read sequencing highlighting the importance of alternative splicing in neuronal development and function^28,30^. The importance of considering coordination between exons is underscored by our observations at the protein level, which reveals adjacent coordinated exons events often create in-frame transcripts that avoid disrupting structured protein domains, likely diversifying and modulating protein function. This observation supports the idea that coordinated splicing events may contribute to the expansion of protein functional diversity without compromising structural protein integrity^6,31^.

The use of direct RNA sequencing allowed us to evaluate the conservation of RNA modifications across evolutionary lineages. Like many RNA processing events^9,18^, m6A modifications display a strong species-specific profile. Nevertheless, a significant subset of m6A sites display conservation across mammals, with these sites often clustering in transcripts close to the stop codon. This clustering pattern is consistent with previous findings on m6A distribution^12,32^ and may suggest a conserved role for these modifications in regulating mRNA stability or translation efficiency^13^. Our findings on the conservation of m6A modifications, particularly those not associated with the canonical DRACH motif, suggest that the deposition of these modifications may be more complex and regulated than previously thought. For example, we identify a substantial number of AG-rich motifs that bear a strong resemblance to the consensus sequence previously identified for METTL16^33^. The higher stoichiometry of AG-rich and non-DRACH m6A sites, despite occurring at lower frequencies, indicates that these sites may have important functional roles that have been conserved across mammalian evolution. These results suggest that m6A deposition is conserved at the single nucleotide level and is driven by more than just the presence of the DRACH motif^24^, supporting a role for other sequence features in determining m6A deposition^34^.

In conclusion, our study provides the first comprehensive long-read overview of the conservation and complexity of mammalian transcriptomes, revealing intricate layers of regulation at the levels of alternative splicing, coordinated splicing, and epitranscriptomic modifications. These findings contribute to our understanding of how transcript complexity has evolved and how it may contribute to phenotypic diversity across mammalian species. Future studies should focus on elucidating the functional consequences of these conserved regulatory mechanisms and their potential roles in development, tissue-specific functions, and disease processes.

## METHODS

### Data reporting

No statistical methods were used to predetermine sample sizes. At least 2 biological replicates were generated for each tissue type across all species. All samples used are listed in Table S1. The experiments were not randomized, and investigators were not blinded to sample allocation during experiments and outcome assessment.

### Sample Collection and ethics statements

The pan-vertebrate dataset comprises six species (human, mouse, rat, dog, cow and chicken) and six tissues (cerebellum, frontal cortex, hippocampus, liver, skeletal muscle and testis). Each tissue had two biological replicates (Table S1). These samples represent the major clades on mammalian phylogeny, with chicken representing an out-group, as well as tissues arising from all the primary germ layers.

Mouse, rat and chicken tissue samples were collected at Garvan Institute of Medical Research and Victor Chang Cardiac Research Institute. Animal use was approved by the Animal Ethics Committee at St. Vincent Hospital, Sydney. Human, dog and bovine total RNA tissues samples (frontal cortex, cerebellum, hippocampus, testis and liver) were obtained from Zyagen, Biochain and Amsbio.

### Tissue dissections

#### Animal maintenance and tissue collection

For mice, 10-12 weeks old C57 Black (C57BL/6J) mice were culled by cervical dislocation followed by decapitation. Brain was isolated and dissected in chilled dissection buffer (1X Hanks’ Balanced Salt Solution [HBSS], 2.5 mM HEPES-KOH [pH 7.4], 35 mM glucose and 4 mM NaHCO_3_) supplemented with fresh 100 μM. Hippocampus, cerebellum, prefrontal cortex, liver, skeletal muscle (thigh) and testis were also isolated, immediately snap frozen with liquid nitrogen and stored at -80°C until use.

Male Long-Evans rats 10 to 12 weeks old were used and anesthetized with an intraperitoneal injection of a mixture of 1.3 mL/kg ketamine (Ketamil 100 mg/mL, Ilium) and 0.3 mL / Kg xylazine (Xylazil 20 mg/mL, Ilium) on the tissue collection day. The whole brain was extracted and placed immediately in a chilled dissection buffer. Similarly to the mice brain preparation described above, the brain was then cut twice coronally using the hypothalamus as the coordinates to expose different compartments. Hippocampus, cerebellum, prefrontal cortex, liver, skeletal muscle (thigh) and testis were also isolated as describe for the mice samples above.

A whole frozen chicken brain was commercially obtained and maintained at -80°C until use. To collect the species equivalent to the cerebellum, frontal cortex and hippocampus, the brain was carefully dissected on a cryostat to expose different brain parts. Tissues were then again snap-frozen with liquid nitrogen until use. Given the anatomical differences between rodent and avian species, prefrontal cortex was collected based on the previously established approaches^35^. As for mouse and rat, hippocampus, cerebellum, liver, skeletal muscle (thigh) and testis were also isolated and stored in the same way.

All research and animal care procedures were approved by the Garvan Institute / St. Vincent’s Hospital Animal Ethics Committee and the University of New South Wales Animal Care and Ethics Committee in accordance with the Australian Code of Practice for the Care and Use of Animals for Scientific Purpose. Mice were housed under conditions of controlled temperature (22°C for standard laboratory temperature) and illumination (12 h light cycle, lights on at 07:00 h). Mice had free access to water and were fed normal chow diet (8 % calories from fat, 21% calories from protein, 71% calories from carbohydrate, 2.6 kcal/g; Gordon’s Speciality Stock Feeds, Yanderra, NSW, Australia).

### RNA extraction

RNA was extracted from the mouse, rat, and chicken tissue samples using a mortar and pestle with liquid nitrogen to reduce the sample size to 50-100 mg. The extracted RNA of each sample was combined with 1 mL of Trizol reagent (Life Technologies, 15596018), followed by tissue disruption on TissueLyzer (Qiagen) using one stainless steel bead of 5 mm (Qiagen cat. no.

69989) per sample at a frequency of 25Hz for 4 minutes. Afterwards, 200 μl of chloroform was added to each sample, mixed by inversion and incubated at room temperature for 2 minutes. After 15 minutes of centrifugation at 12,000g at 4°C, the upper aqueous phase was collected and transferred to a new 1.5 ml microcentrifuge tube (Eppendorf). RNA samples were precipitated by adding 500 μl of isopropanol followed by overnight incubation at -20°C and centrifugation at 7,500 g for 15 minutes at 4°C. The supernatants were then discarded, and RNA pellets were washed with 75% (v/v) ethanol and centrifuged at 7,500 g for 5 minutes at 4°C.

Supernatants were discarded, and pellets air dried at room temperature. RNA pellets were then dissolved in 50 μl nuclease-free water with brief heatingto 65°C. DNAse treatment was performed by adding 3 μL of TURBO DNase (Ambion, cat. no. AM2239) per 30 μg of total RNA per sample and incubating for 40 minutes at 37°C. Samples were then purified by overnight ethanol precipitation (2.5 volumes of 100% ethanol and 0.1 volume of 3M NaOAc pH 5.2; see above for the precipitation and collection procedure) at -20°C or column clean up using RNeasy MinElute Cleanup Kit (Qiagen, cat. no. 74204).

### RNA quality analysis

The RNA purity and concentration were measured using a NanoDrop spectrophotometer (Thermo Fisher Scientific), and RNA integrity was verified on Agilent 4200 TapeStation with RNA screen Tape (Table S1). The Agilent 4200 TapeStation reports the percentage of RNA fragments > 200nt (DV200 values), which estimate the quality of the RNA samples before sequencing. To ensure that our samples had long RNA molecules for Nanopore direct RNA sequencing, we also calculated the percentage of RNA fragments > 500 nucleotides (DV500) by manually selecting in the TapeStation Software the cutoff of the distribution equal to 500 nucleotides.

### Nanopore direct RNA sequencing

#### RNA library preparation

Total RNA libraries for direct RNA sequencing (DRS) were prepared generally according to the manufacturer’s protocol provided by Oxford Nanopore Technologies (ONT), with minor deviations for optimisation purposes. About 5,000 ng of RNA from each tissue was used for sequencing with MinION and PromethION flowcells, employing the SQK002 direct RNA sequencing kit. Briefly, the protocol consists of six steps. Initially, the RNA sample underwent sequential mixing with T4 DNA Ligase Buffer (Thermo Fisher Scientific), RT Adapter RTA, T4 DNA Ligase HC, and RNassin Plus (Promega) for the ligation of RT Adaptor, followed by an incubation period as prescribed by the ONT SQK002 protocol. Subsequently, reverse transcription was carried out by supplementing the mixture with 10 mM dNTPs, 5× SS UV First Strand Buffer, 0.1M DTT, Nuclease-free water, and Superscript IV RT. The resulting library underwent purification using Agencourt AMPure XP beads (Beckman), involving incubation, magnetic pelleting, ethanol washing, and elution steps. In the subsequent ligation step, the purified library was combined with RNA Adaptor RMX, T4 DNA Ligase HC, T4 DNA Ligase buffer, and RNA-grade deionised water to facilitate ligation of the library’s fragments with a motor adaptor as per manufacturer’s instructions. A final cleanup procedure using Agencourt AMPure XP beads was then performed, followed by elution in elution buffer ELB. All incubation durations, concentrations, and temperatures were rigorously maintained in accordance with the SQK002 protocol provided by ONT. The resulting RNA library was subsequently directly used for loading or preserved at 4°C prior to loading.

### Flow cell loading and live sequencing setup

The library loading procedure adhered to the protocol outlined by ONT for the MinION Mk1B or PromethION platform. The flow cell was inserted under the clip of the MinION, ensuring proper thermal and electrical contact. Quality control of the flow cell was performed, and the priming port was opened to remove any air. A priming mix, comprising Flush Tether and Flush Buffer (Activated Flush Buffer), was prepared, degassed, and loaded into the flow cell as per the manufacturer’s instructions. Following this, the RNA library was prepared by mixing RNA Running Buffer, the ELB-eluted RNA and RNA-grade deionised water. Upon degassing, the library was gently loaded into the flow cell *via* the SpotON sample port in a dropwise manner. Finally, the priming port was closed, the SpotON sample port cover was replaced, and the MinION or PromethION was secured in place, following the recommended protocol guidelines. The DRS sequencing runs were conducted at room ambient temperature (∼25°C) up to a maximum duration of 72 hours. Live basecalling was disabled during the run and the file format for data acquisition was set as FAST5 files for downstream preprocessing steps and subsequent bioinformatics analysis.

### Construction of orthologous transcript dataset

To overcome discrepancies in the annotation quality of species sequenced in this study within public databases, we supplemented the annotations from Ensembl (release version 102) with human annotations, since human is the best-annotated species. This aimed to ensure differences between organisms were due to biological differences and not annotation biases.

Ensembl Biomart^36^ was used to obtain 1-to-1 orthologues (orthologues that come from a speciation event) for all human protein-coding genes between the six species. To obtain the orthologous region of the exons forming each transcript, every exon in orthologous transcripts was mapped over from human to each species using LiftOver (with parameter -minMatch = 0.1) and UCSC chain files^37^. Lifted over regions overlapping annotated exons were corrected to match the closest annotated splice sites (Ensembl 102 annotation) within 100nt. If the mapped over region did not match an annotated exon, the 3’ and 5’ splice sites were corrected to nearest annotated AG and GT residues, respectively. Lifted over regions outside orthologous gene bodies were not included in the analysis. The orthologous transcripts were then reassembled into transcripts using the exon order from annotated human transcripts. These transcripts were then annotated with original human transcript ID to facilitate correspondence for future analysis. Transcripts conserved across all studied species were considered as “genomic conserved transcripts”.

To ensure species-specific transcripts were also included in the annotation, transcripts from Ensembl annotation (release version 102) from non-1-to-1 orthologues (Table S2) and not identified as genomic conserved transcripts (Table S3) were included in the annotation. These data were collated in GTF files for further analysis (Supplementary Data Files).

### Nanopore base calling, read mapping, and transcript quantification

The nanopore data was processed using guppy version 5.0.7+2332e8d to basecall the nucleotide sequences. The parameters used in guppy were: --flowcell FLO-MIN106 --kit SQK-RNA002 --device cuda:all:100% --compress_fastq --recursive --num_callers 4 -- gpu_runners_per_device 2 --chunks_per_runner 512 --chunk_size 1000. The total number of reads per run is given in Table S1.

To extract the transcriptome fasta sequence for each species, the program gffread (https://github.com/gpertea/gffread) in combination with our supplemented GTF files was used. In the case of human, we filtered the GTF file (release version 102) to contain protein-coding transcripts, and we built the transcriptome file using gffread. After basecalling and mapping, we performed a quality analysis of total reads and of mapped reads using Nanoplot (v1.41.0) and NanoComp (De Coster and Rademakers 2023). The parameters used were --fastq for fastq files and --bam for mapped reads (bam files input).

Nanopore read sequences were mapped to the transcriptome of each species using minimap2 version 2.24-r1122^38^ using parameters -ax map-ont -N 10. The percentage of mapped reads (mapping percentage) was calculated to assess the quality of the transcriptomes with extra orthologue transcripts (Figure S1B) (Table S1). The mapped reads were those without secondary or supplementary alignments according to samtools^39^. The command used in samtools was: samtools view -F 0×904 -c <file.sorted.bam>. After mapping, NanoCount v1.1.0^40^ was used to quantify the expression of the transcripts, using default parameters together with -- extra_tx_info. The percentage of transcript expression per gene (PTE) was calculated as a percentage of the total gene expression (*i*.*e*. the sum of all transcript expression for a gene). This was done for each sample and the average across replicates was used when stated.

### Assessment of full-length read mapping *vs*. total reads mapping

To validate if the sequencing reads obtained by direct RNA-sequencing were full-length, we used the script from Gleeson et al. 2022^40^, BamSlam (https://github.com/josiegleeson/BamSlam) to identify the full-length transcript isoforms. Briefly, we used the bam output files from NanoCount of hippocampus samples of human and input to BamSlam. Then, BamSlam calculates the coverage fractions by dividing the alignment length by the original known transcript length of each read’s best alignment. We used two cutoffs to define full-length reads: 90% and 95% of coverage in the annotated transcript. Afterwards, we subset the full-length reads and re-mapped them to the transcriptome. To assess if truncated reads *vs*. full-length reads had a disproportionate mapping to a region of the transcript (bias towards the 3’ vs 5’ region), we calculated the position of exons mapped to 100 percentile bins. For this, we define *exonchunks* of overlapping exons of each transcript of the same gene and assign each exon to a bin. We then calculated the percentage of exons in each bin. We performed such calculations on all the reads, 90% of coverage reads, and 95% of coverage reads.

### Gene and transcript expression clustering analysis

Using NanoCount transcript expression, we calculated gene expression for all orthologue genes and transcripts. Only transcripts with at least CPM > 1 and total gene expression CPM > 3 in two samples across species were included in the analysis. We performed UMAP clustering for transcripts across all four quartiles of relative gene expression using the umap R package (N-components = 2, random_state = 5000, n_neighbours = 24). Relative expression was calculated as the fraction of transcript contributing to total gene expression on a tissue and was the equivalent to the percent spliced in (PSI) used in exon splicing analysis. A transcript in the lowest quartile represents transcripts contributing a maximum of 25% of total gene expression.

### Identification of conserved, tissue-specific, and switching transcripts

Transcripts are only defined as expressed in a species or a tissue if they have CPM > 1. These transcripts can have unique exonic structures or be “genomically conserved transcripts”. Transcripts are genomically conserved if their exon structure is conserved across species (*i*.*e*. these were included in our custom GTF files). A gene with multiple transcripts must therefore have at least two transcripts with CPM > 1 in the same species. A gene with multiple conserved transcripts must have at least two genomically conserved transcripts with CPM > 1 in the same species. A “mammalian conserved transcript” must therefore have CPM > 1 in at least one tissue in all mammalian species analysed. A “vertebrate conserved transcript” must also be expressed with CPM > 1 in chicken as well. Switching transcripts are defined as those cases where the major transcript isoform (defined as the transcript in a gene with the largest PTE) is different among tissue samples. A tissue-specific transcript is the major transcript of a gene in a single tissue.

To calculate alternative exon usage, a custom GTF annotation was constructed for all genes with multiple conserved transcripts. SUPPA2^41^ subcommand “generateEvents” with default settings was then used to generate transcript events: Skipping Exon (SE), Alternative 5’/3’ Splice Sites (A5/A3), Mutually Exclusive Exons (MX), Retained Intron (RI), Alternative First/Last Exons (AF/AL). The proportion of these events was compared between categories and plotted using ggplot.

### Coordinated splicing analysis

The coordinated splicing analysis uses previously published approach (available at https://github.com/pgupta3005/Coordinated_splicing). Briefly, this pipeline requires a GTF file and a counts table as input. The GTF is used to create a database of transcript and exon information ensuring unique entries for each exon, so exons can be compared across transcripts. Overlapping exons irrespetive of their start and end coordinates are merged as the same exons. Next, a set of “expressed exons” is defined by removing genes with only one transcript and genes with less than ten counts. This operation removes exons present in only 5% of the gene transcripts, as well asmono- and di-exon genes. The pipeline next identifies transcripts that match all splice junctions (Full Spliced Match, FSM, (Tardaguila et al. 2018)) and transcripts that are shorter versions (fragmented) of the FSM transcript defining the transcription start sites (TSS) and transcription termination sites (TTS) for each transcript. Next, constitutive exon chunks are defined as exons with greater than 95% inclusion across all gene transcripts. Thus, alternative exon chunks have less than 95% inclusion in a gene’s transcript. Then, the pipeline defines pairs of exons, the categories of coordinated splicing events and their absolute positions (location of the exons within the gene). The relative position of the exon chunks is split into two categories: (1) adjacent – two consecutive alternative exons, (2) distant – with one or more constitutive exons in between. These relative positions define the type of coordinated splicing events (Figure 2A). Then, the pipeline also defines the absolute positions (start, internal and end) of each exon chunk according to the gene body. Finally, the pipeline calculates whether the exon coordinate chunks show greater coordination than by random chance. For this, counts of transcripts with coordinated exons are identified to make a contingency table and calculate the p-value and log odds ratio (LOR) for the Fisher’s test. A false discovery rate (FDR) is calculated for all exon chunks defined as adjacent and distant events. The sign of the LOR value defines if an event is mutually associated (exons are always present in a transcript) or mutually exclusive (only one exon is present in a transcript). The coordinated splicing pipeline was run for each of the six tissues of the human dataset.

### Validation of coordinated splicing with full length reads

As we have previously described, full-length reads were obtained using the BamSlam script from Gleeson et al. 2022^40^. These reads were re-mapped to the transcriptome, and NanoCount was used to obtain transcript expression data. Then, the coordinated splicing pipeline was run with counts expression data of 90% and 95% full-length reads.

### Exon variability

Exon variability was defined as previously described in Hardwick et al. (2022)^22^. Briefly, we defined a set of conserved exons for each of the tissues evaluated for both adjacent and distant coordinated splicing events. For each set of variable exons maximum and minimum PSI values across tissues was calculated. The log_2_ fold difference between maximum and minimum PSI values was plotted using R package ggplot.

### Conserved and non-conserved coordinated splicing events

Coordinated splicing events were defined as conserved or non-conserved if the exon-chunks forming the event were present in conserved or non-conserved transcripts. For a coordinated splicing event to be defined as conserved, both exon-chunks must be present in a conserved transcript. Similarly, for a coordinated splicing event to be defined as non-conserved, at least one of the exon-chunks must be present in a non-conserved transcript. The transcript’s conservation was defined in terms of exons that have an orthologous region to human exons of the same transcript.

### Protein analysis

Events retaining protein sequence were defined as exons that were divisible by 3, lack in-frame stop codon and exons that either the downstream or upstream exon are a UTR-encoding exon. For all positions in a protein a score for intrinsic disorder was computed using IUPred (iupred.enzim.hu)^42^. Amino acid residues with a score greater than 0.4 were considered disordered. For each coding exon the fraction of disordered residues was estimated. Regions within structured domains were defined using UniProt^43^ and Pfam^44^ annotation.

### Gene Ontology enrichment analysis of genes with coordinated splicing events in more than one tissue

Gene Set Enrichment analysis was performed using either the goseq() R function from the multiGSEA R package^45^ or gProfiler. For multiGSE the Gene Ontology (GO) database was retrieved using the getMSigGeneSetDb() function in R with the parameters: species = “human” and id.type = “ensembl”. For gProfiler, default datasets for Gene Ontology were used and default settings with a maximum gene set of 2000. The background dataset used was composed of orthologue genes identified across all six species from our pan-vertebrate dataset expressed in human samples (> 1 CPM in at least 2 samples). The output of the GO enrichment analysis was filtered according to a False Discovery Rate cutoff of 0.05. The resulting terms were plotted using the ggplot() function from the ggplot2 library in R.

### RNA m6A modification detection

All reads from the sequencing runs for each species and tissues were pooled together for the m6A analysis. For each species and tissue, we run CHEUI^15^ on the reads mapped to the corresponding transcriptomes defined above, filtered to include the two most highly expressed transcripts per a tissue and longest expressed transcripts per a gene. CHEUI predicts m6A using the nanopore signals corresponding to a 9-mer context for every A-site, *i*.*e*. NNNN<u>A</u>NNNN. We run model 1 to predict m6A in individual reads at every possible A-site in the expanded transcriptome for each species. We then used CHEUI model 2 to predict m6A at all transcriptomic A-sites with >20 reads.

We tested an average of 3.5 million transcriptomic A sites in each condition, considering all sequence contexts (Table S6). As previously described^15^, the false positive rate was estimated by performing a permutation test of the individual read modification probabilities across sites, maintaining the same read coverage per transcript site (script provided in https://github.com/comprna/CHEUI). We selected a probability score cutoff of P>0.998 based on an estimated false positive rate of 0.0000-0.0003 across samples after performing a permutation of the reads for all sites maintaining the coverage per site. This resulted in an average of 27,000 sites per tissue and species (Table S6).

### Mapping and annotating m6A modifications in genomic coordinates

Tested and significant (P>0.998) m6A positions in the transcriptome were mapped onto the genomic coordinates of the corresponding species using R2Dtool^46^, using the transcriptome-based bed file of the m6A sites and the extended GTF file from each species described above. We also used R2Dtool to annotate positions of the tested and significant sites relative to the start and stop codons mRNAs and exon-exon junctions and to generate the modification density profiles. Coordinates for all tested and significant sites in each species and tissue were then mapped to human assembly using the CrossMap^47^, achieving a mapping efficiency of >95% across mammals and about 75% for chicken (Table S7). When multiple significant transcriptomic sites mapped to the same genomic site, we selected the stoichiometry and probability of the transcriptomic site with the highest probability.

### Conservation of m6A modifications

The conservation of human m6A sites was estimated using adenosine sites that were also tested in other species. We then calculated the proportion of human m6A sites that were pairwise conserved in the other species, as well as the proportion of human m6A sites that were conserved in one or more species, two or more species, *etc*. In all those cases, the conservation in a given species was conditioned to the site being tested in that species. For the heatmap in Figure 3, only adenosines with at least one m6A modification in any tissue and that were tested in every species were considered. From this dataset, we considered as “mammalian conserved” the human adenosines that were tested across all mammals and found significant in human and two or more other mammals. We also considered as “human-specific” the human m6A sites that were tested in other species but were found not to be significant in those other species.

### Motiif contexts of the m6A modifications

The motifs were defined using regular expressions against the 9-mer sequences identified by CHEUI with the central residue being m6A modified. The DRACH motif was defined by the regular expression “..[AGT][GA]AC[ACT]..”. The DRACH-like motif was defined by “…[GA]AC[ACT]..|..[AGT][GA]AC…”, which is equivalent to a DRAC and RACH motifs. The AG-rich motif was defined by “A[AG][AG][AG].” The non-DRACH was defined as 9-mers that did not match any of the above regular expressions.

### Conservation of the m6A sequence context

For each m6A site, either mammalian conserved or human-specific, we considered the 5-mer centred on the m6A site. We used the base-wise phyloP score from the multiple sequence alignments of human with 100 vertebrates (phyloP100way).

### Isoform specificity of the m6A modifications

3’UTRs differentially regulated through alternative polyadenylation were identified from expressed transcripts (count > 5) across all analysed human tissues. The minimum and maximum coordinates of the end of the 3’UTR for each gene were identified. An m6A modification was considered as differentially regulated by alternative polyadenylation, if it occurred between the minimum and maximum coordinates and therefore was located in the alternative sequence differentially included between transcripts of the same gene.

### ClinVar and COSMIC analysis

To identify m6A modifications overlapping putative disease mutations, we downloaded data from ClinVar (clinvar_20240730.vcf.gz)^48^ and COSMIC (08/2024)^49^ databases. ClinVar data was filtered to remove all variants annotated as benign and only point mutations from COSMIC were considered. To account for the known sequence specificity of m6A modifications, the nucleotides immediately upstream and downstream of m6A site were all taken into consideration. Both the filtered ClinVar and COSMIC datasets, as well as the m6A sites, were converted into bed files and the command bedtools intersect (default settings) was used to determine the overlaps.

### Statistical analyses

Statistical analyses were done using R programming language version 4.3.1 (2023-06-16). Distributions of variables across the samples were compared using the Wilcoxon rank sum test with the function wilcox.test(). Fisher exact test was used to assess whether two categorical variables were independent. For these analyses, the oddsratio() function from the epitools R package was used. The input for the oddsratio function is a contingency table of the frequency of each variable. For multiple test correction, the p.adjust() function with the Benjamin and Hochberg correction (parameter “BH”) was used. For data visualisation, outliers with a lower limit as Q1-1.5*IQR and an upper limit as Q3+1.5*IQR were removed.

## DATA AVAILABILITY

Raw and processed direct nanopore long read RNA-seq data have been deposited in SRA.

## CODE AVAILABIILTY

Code is available for coordinated splicing analysis at:

https://github.com/pgupta3005/Coordinated_splicing and m6A analysis at: https://github.com/comprna/CHEUI

