## Supplementary Figures for "The conserved landscape of RNA modifications and transcript diversity across mammalian evolution"

### Slide 1
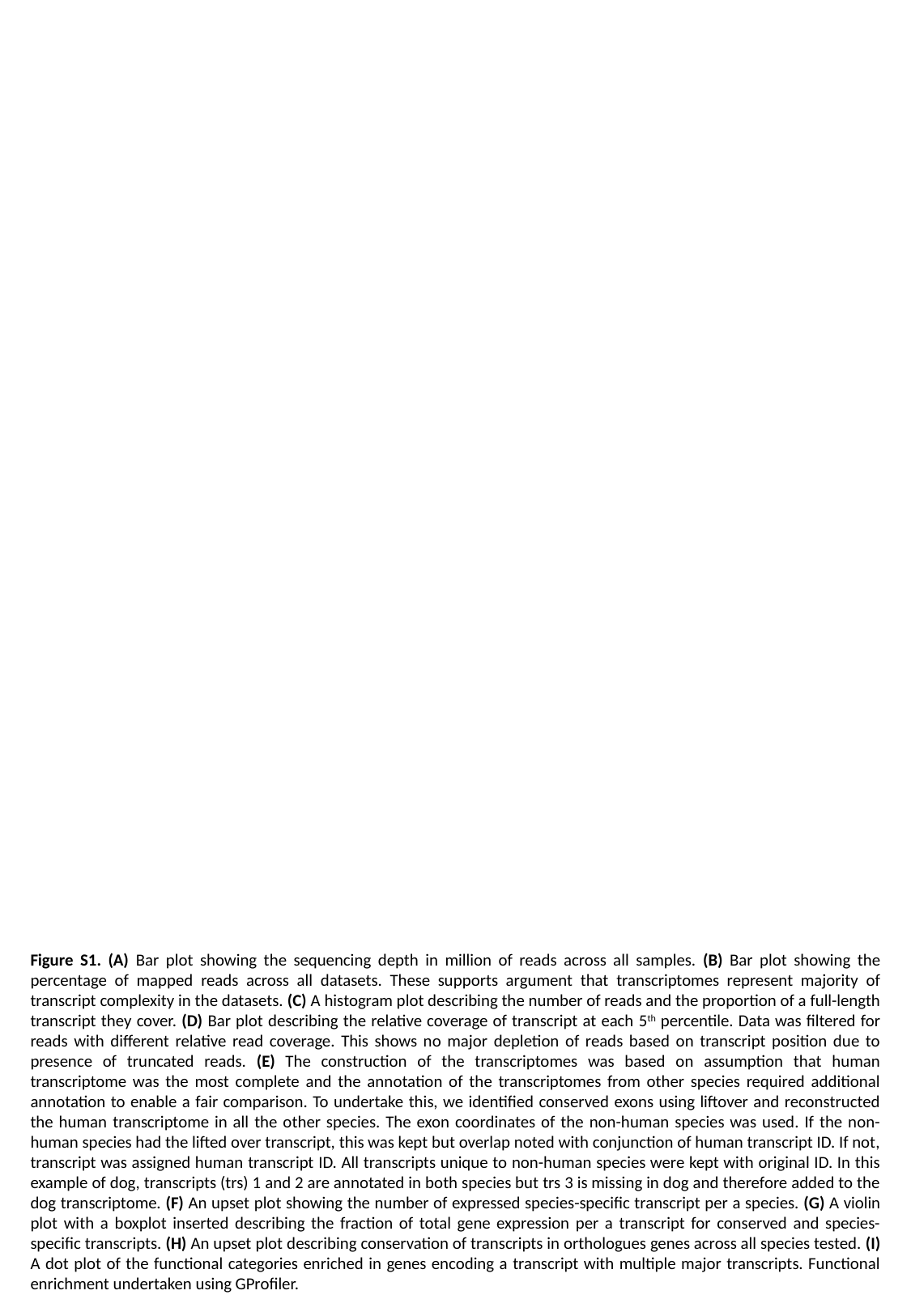

Figure S1. (A) Bar plot showing the sequencing depth in million of reads across all samples. (B) Bar plot showing the percentage of mapped reads across all datasets. These supports argument that transcriptomes represent majority of transcript complexity in the datasets. (C) A histogram plot describing the number of reads and the proportion of a full-length transcript they cover. (D) Bar plot describing the relative coverage of transcript at each 5^th^ percentile. Data was filtered for reads with different relative read coverage. This shows no major depletion of reads based on transcript position due to presence of truncated reads. (E) The construction of the transcriptomes was based on assumption that human transcriptome was the most complete and the annotation of the transcriptomes from other species required additional annotation to enable a fair comparison. To undertake this, we identified conserved exons using liftover and reconstructed the human transcriptome in all the other species. The exon coordinates of the non-human species was used. If the non-human species had the lifted over transcript, this was kept but overlap noted with conjunction of human transcript ID. If not, transcript was assigned human transcript ID. All transcripts unique to non-human species were kept with original ID. In this example of dog, transcripts (trs) 1 and 2 are annotated in both species but trs 3 is missing in dog and therefore added to the dog transcriptome. (F) An upset plot showing the number of expressed species-specific transcript per a species. (G) A violin plot with a boxplot inserted describing the fraction of total gene expression per a transcript for conserved and species-specific transcripts. (H) An upset plot describing conservation of transcripts in orthologues genes across all species tested. (I) A dot plot of the functional categories enriched in genes encoding a transcript with multiple major transcripts. Functional enrichment undertaken using GProfiler.

### Slide 2
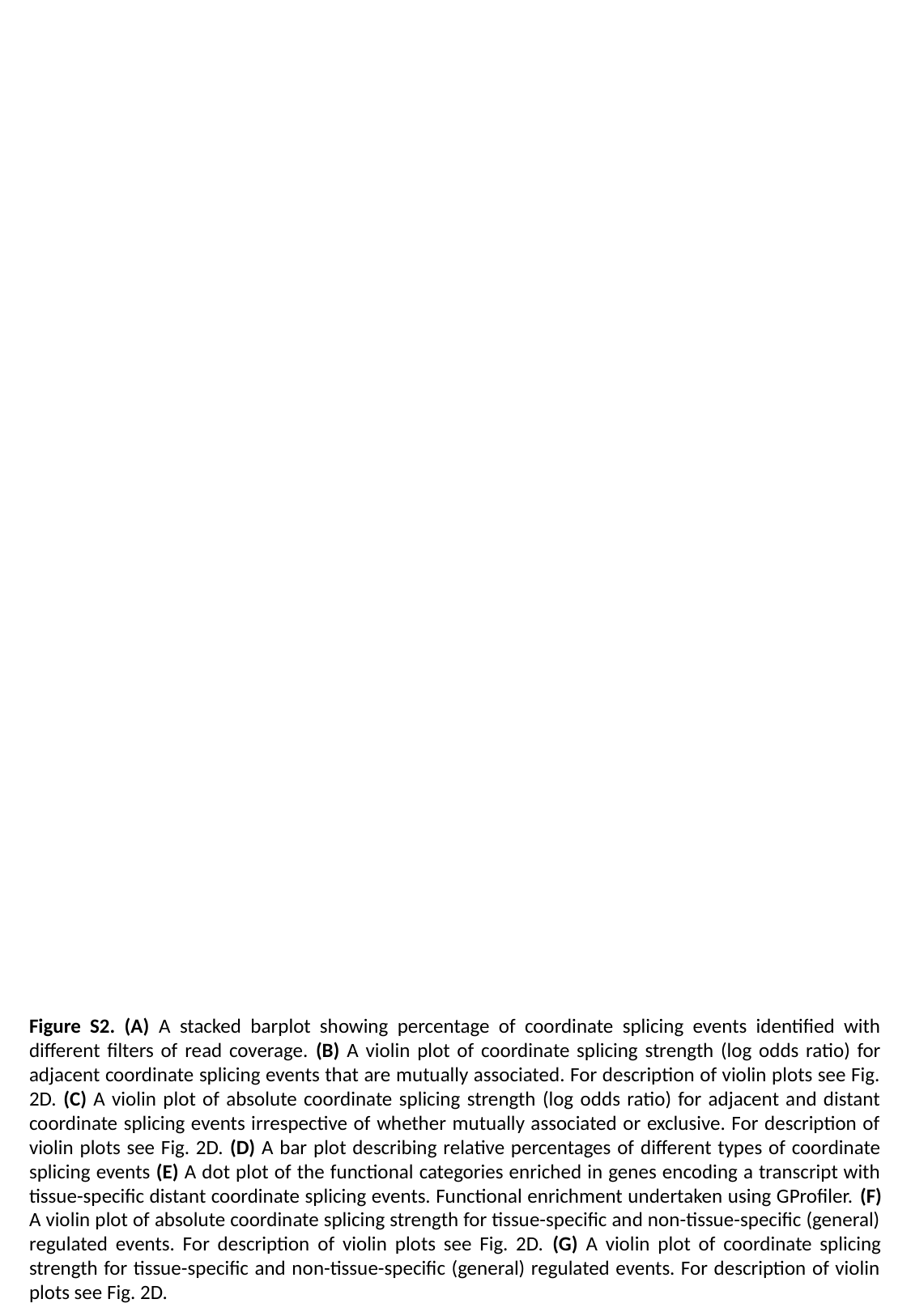

Figure S2. (A) A stacked barplot showing percentage of coordinate splicing events identified with different filters of read coverage. (B) A violin plot of coordinate splicing strength (log odds ratio) for adjacent coordinate splicing events that are mutually associated. For description of violin plots see Fig. 2D. (C) A violin plot of absolute coordinate splicing strength (log odds ratio) for adjacent and distant coordinate splicing events irrespective of whether mutually associated or exclusive. For description of violin plots see Fig. 2D. (D) A bar plot describing relative percentages of different types of coordinate splicing events (E) A dot plot of the functional categories enriched in genes encoding a transcript with tissue-specific distant coordinate splicing events. Functional enrichment undertaken using GProfiler. (F) A violin plot of absolute coordinate splicing strength for tissue-specific and non-tissue-specific (general) regulated events. For description of violin plots see Fig. 2D. (G) A violin plot of coordinate splicing strength for tissue-specific and non-tissue-specific (general) regulated events. For description of violin plots see Fig. 2D.

### Slide 3
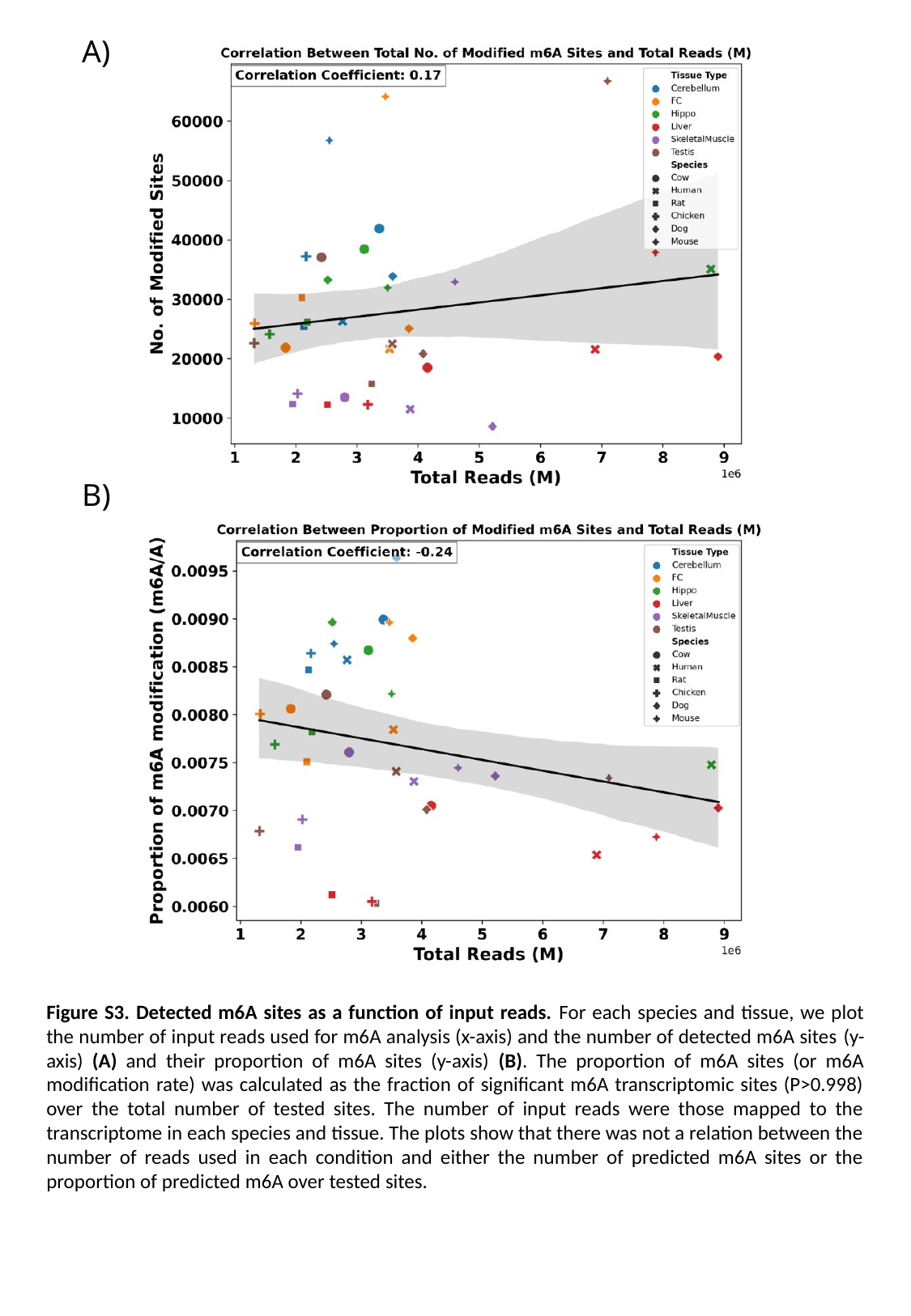

A)
B)
Figure S3. Detected m6A sites as a function of input reads. For each species and tissue, we plot the number of input reads used for m6A analysis (x-axis) and the number of detected m6A sites (y-axis) (A) and their proportion of m6A sites (y-axis) (B). The proportion of m6A sites (or m6A modification rate) was calculated as the fraction of significant m6A transcriptomic sites (P>0.998) over the total number of tested sites. The number of input reads were those mapped to the transcriptome in each species and tissue. The plots show that there was not a relation between the number of reads used in each condition and either the number of predicted m6A sites or the proportion of predicted m6A over tested sites.

### Slide 4
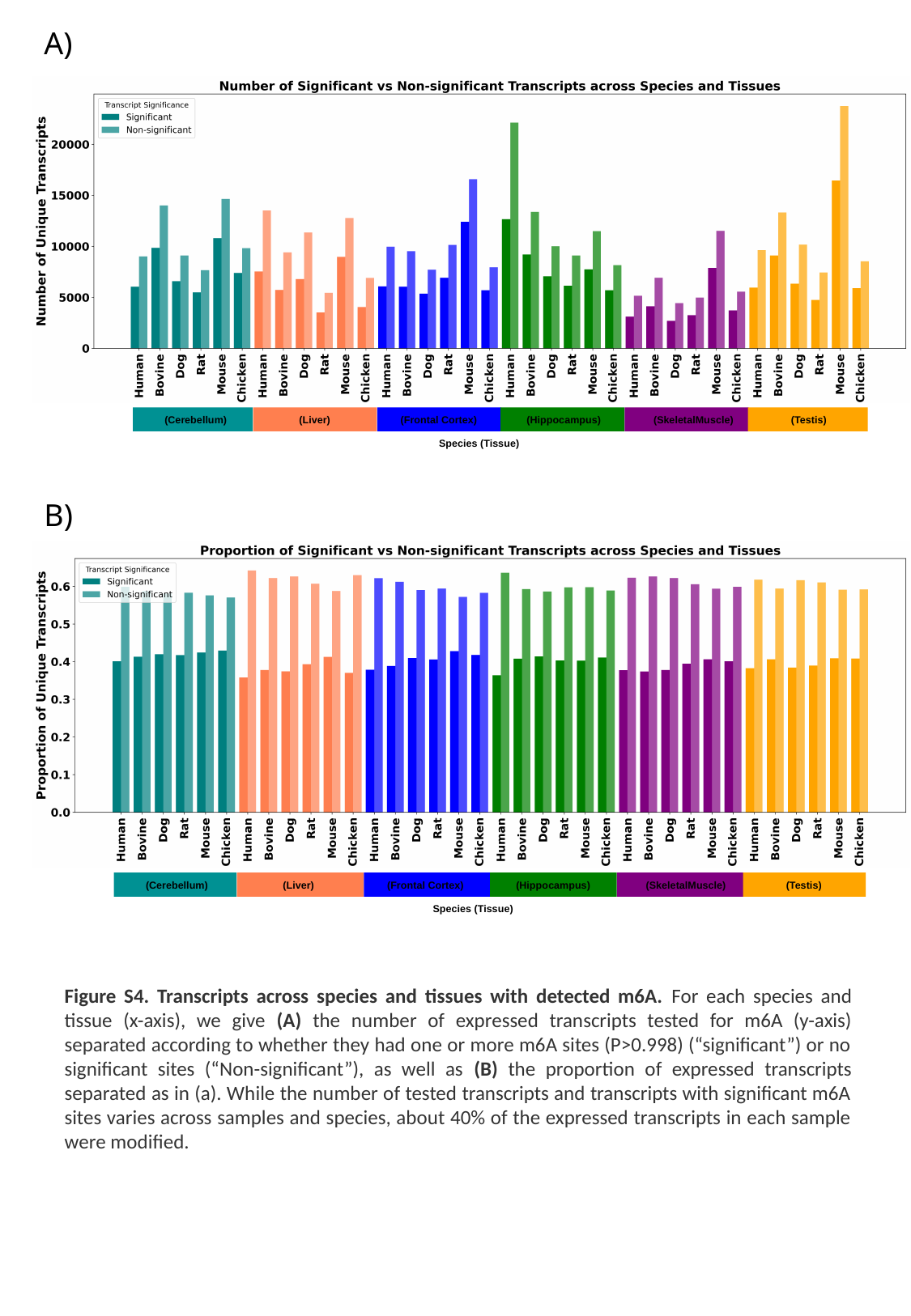

A)
 (Cerebellum)
 (Liver)
 (Frontal Cortex)
 (Hippocampus)
 (SkeletalMuscle)
 (Testis)
Species (Tissue)
B)
 (Cerebellum)
 (Liver)
 (Frontal Cortex)
 (Hippocampus)
 (SkeletalMuscle)
 (Testis)
Species (Tissue)
Figure S4. Transcripts across species and tissues with detected m6A. For each species and tissue (x-axis), we give (A) the number of expressed transcripts tested for m6A (y-axis) separated according to whether they had one or more m6A sites (P>0.998) (“significant”) or no significant sites (“Non-significant”), as well as (B) the proportion of expressed transcripts separated as in (a). While the number of tested transcripts and transcripts with significant m6A sites varies across samples and species, about 40% of the expressed transcripts in each sample were modified.

### Slide 5
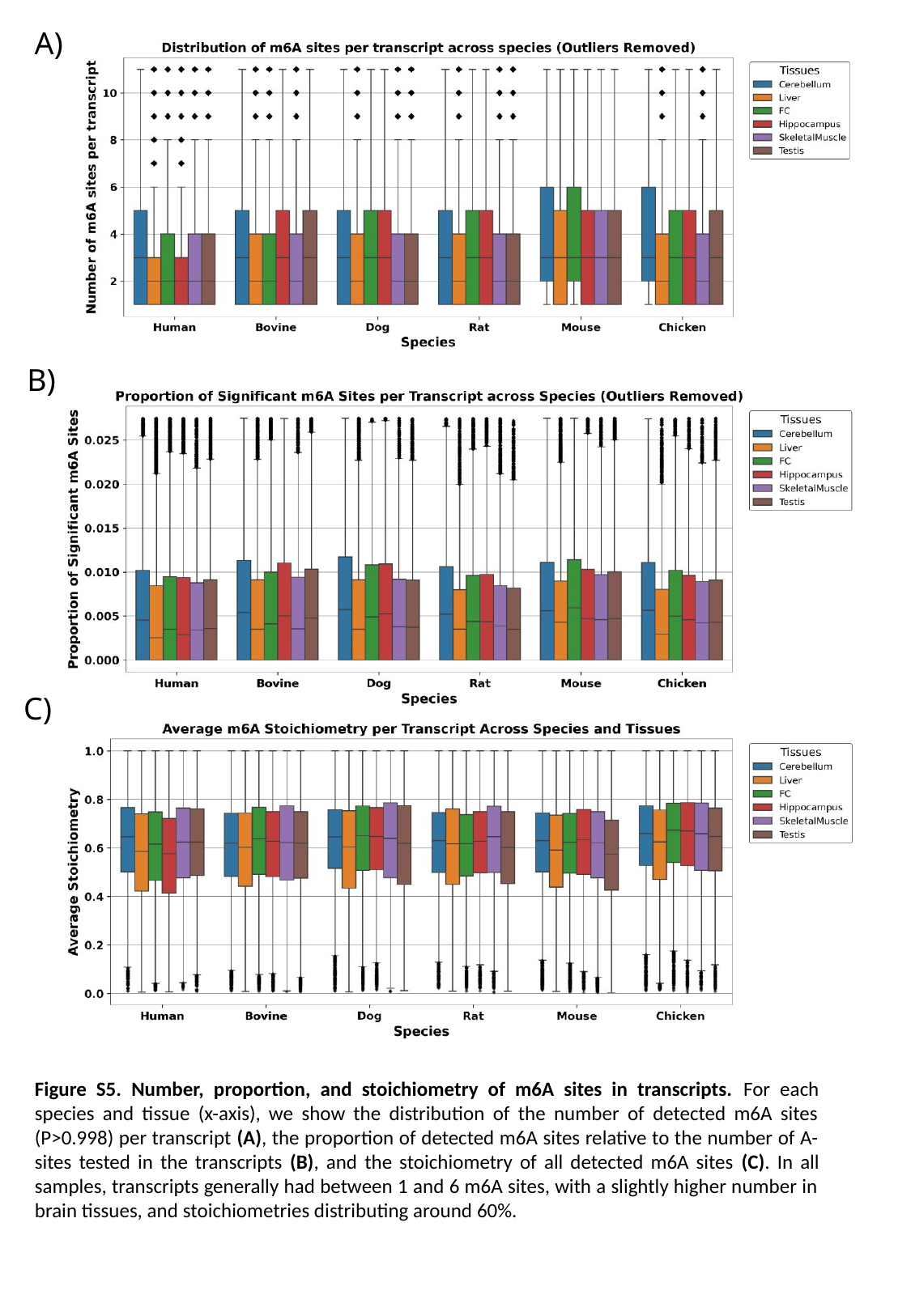

A)
B)
C)
Figure S5. Number, proportion, and stoichiometry of m6A sites in transcripts. For each species and tissue (x-axis), we show the distribution of the number of detected m6A sites (P>0.998) per transcript (A), the proportion of detected m6A sites relative to the number of A-sites tested in the transcripts (B), and the stoichiometry of all detected m6A sites (C). In all samples, transcripts generally had between 1 and 6 m6A sites, with a slightly higher number in brain tissues, and stoichiometries distributing around 60%.

### Slide 6
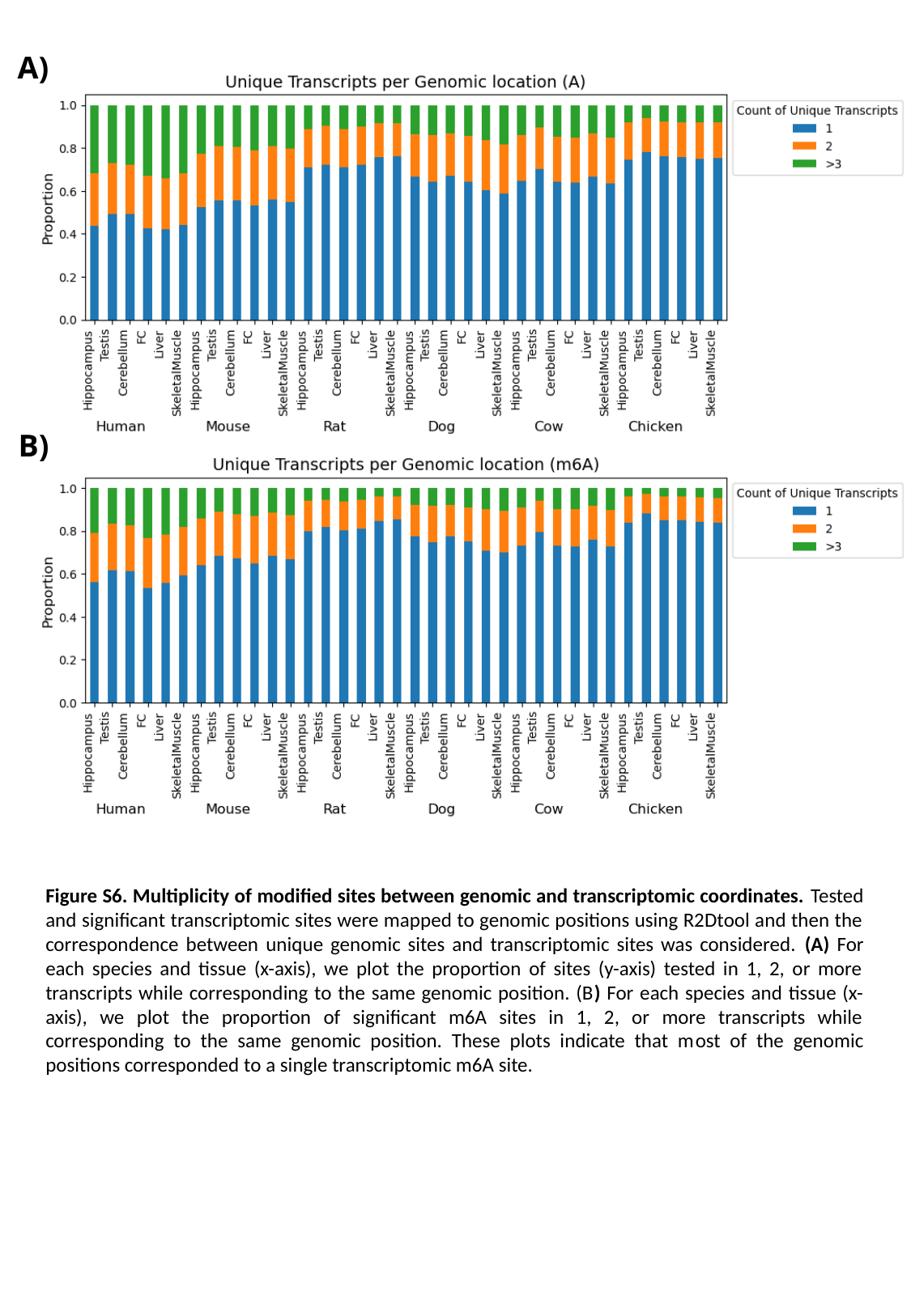

A)
B)
Figure S6. Multiplicity of modified sites between genomic and transcriptomic coordinates. Tested and significant transcriptomic sites were mapped to genomic positions using R2Dtool and then the correspondence between unique genomic sites and transcriptomic sites was considered. (A) For each species and tissue (x-axis), we plot the proportion of sites (y-axis) tested in 1, 2, or more transcripts while corresponding to the same genomic position. (B) For each species and tissue (x-axis), we plot the proportion of significant m6A sites in 1, 2, or more transcripts while corresponding to the same genomic position. These plots indicate that most of the genomic positions corresponded to a single transcriptomic m6A site.

### Slide 7
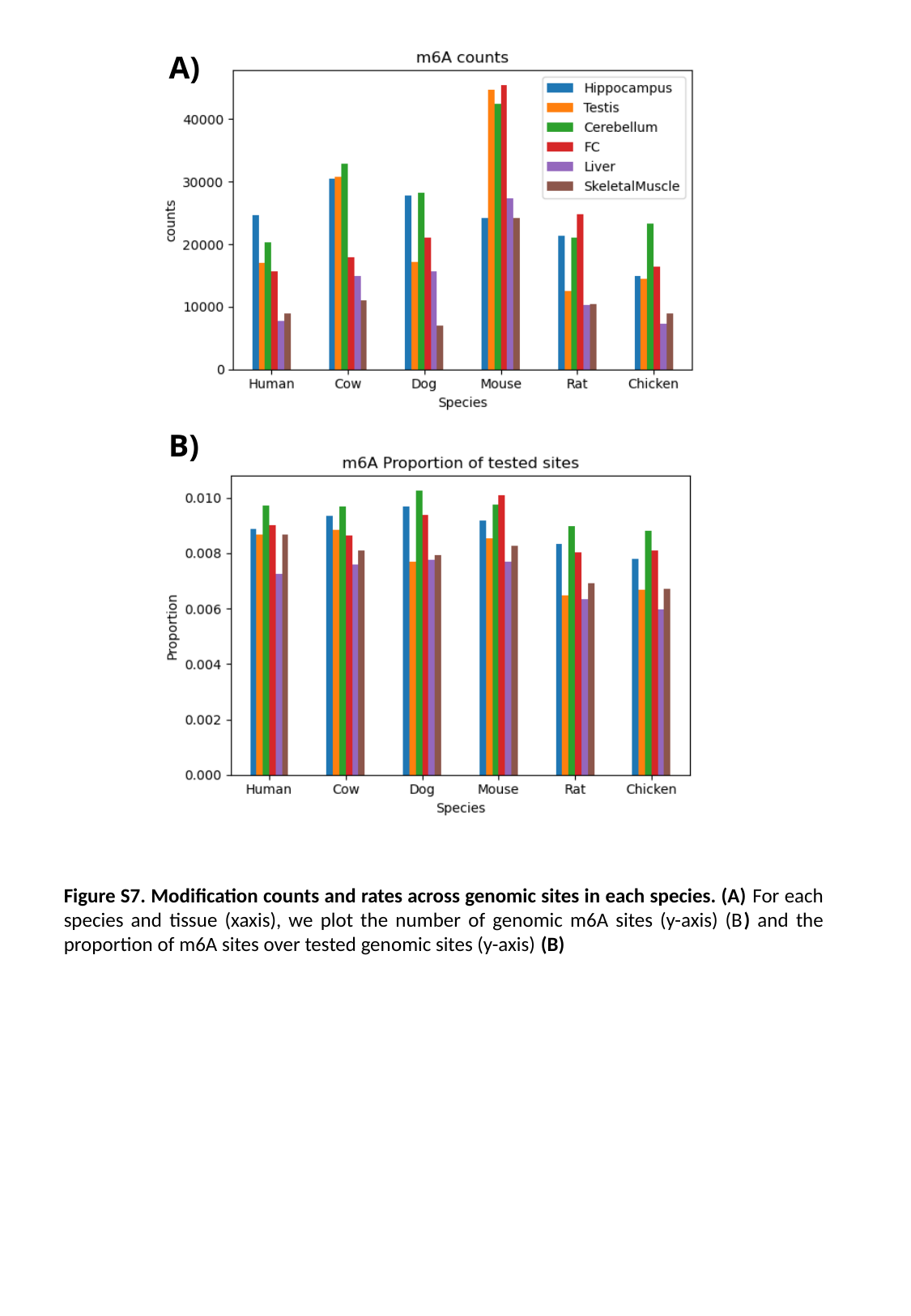

A)
B)
Figure S7. Modification counts and rates across genomic sites in each species. (A) For each species and tissue (xaxis), we plot the number of genomic m6A sites (y-axis) (B) and the proportion of m6A sites over tested genomic sites (y-axis) (B)

### Slide 8
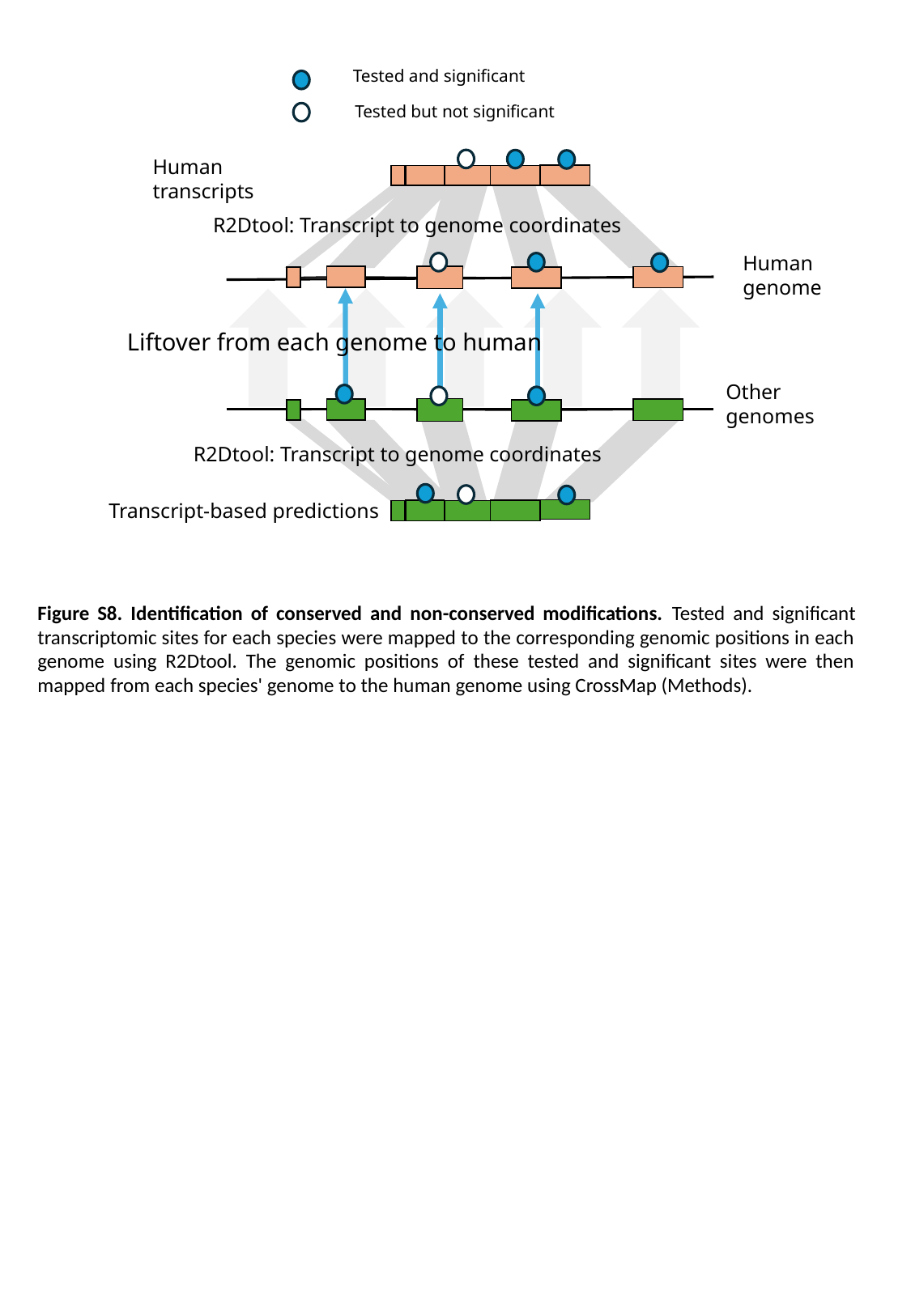

Tested and significant
Tested but not significant
Human transcripts
Human
genome
Liftover from each genome to human
Other
genomes
R2Dtool: Transcript to genome coordinates
Transcript-based predictions
R2Dtool: Transcript to genome coordinates
Figure S8. Identification of conserved and non-conserved modifications. Tested and significant transcriptomic sites for each species were mapped to the corresponding genomic positions in each genome using R2Dtool. The genomic positions of these tested and significant sites were then mapped from each species' genome to the human genome using CrossMap (Methods).

### Slide 9
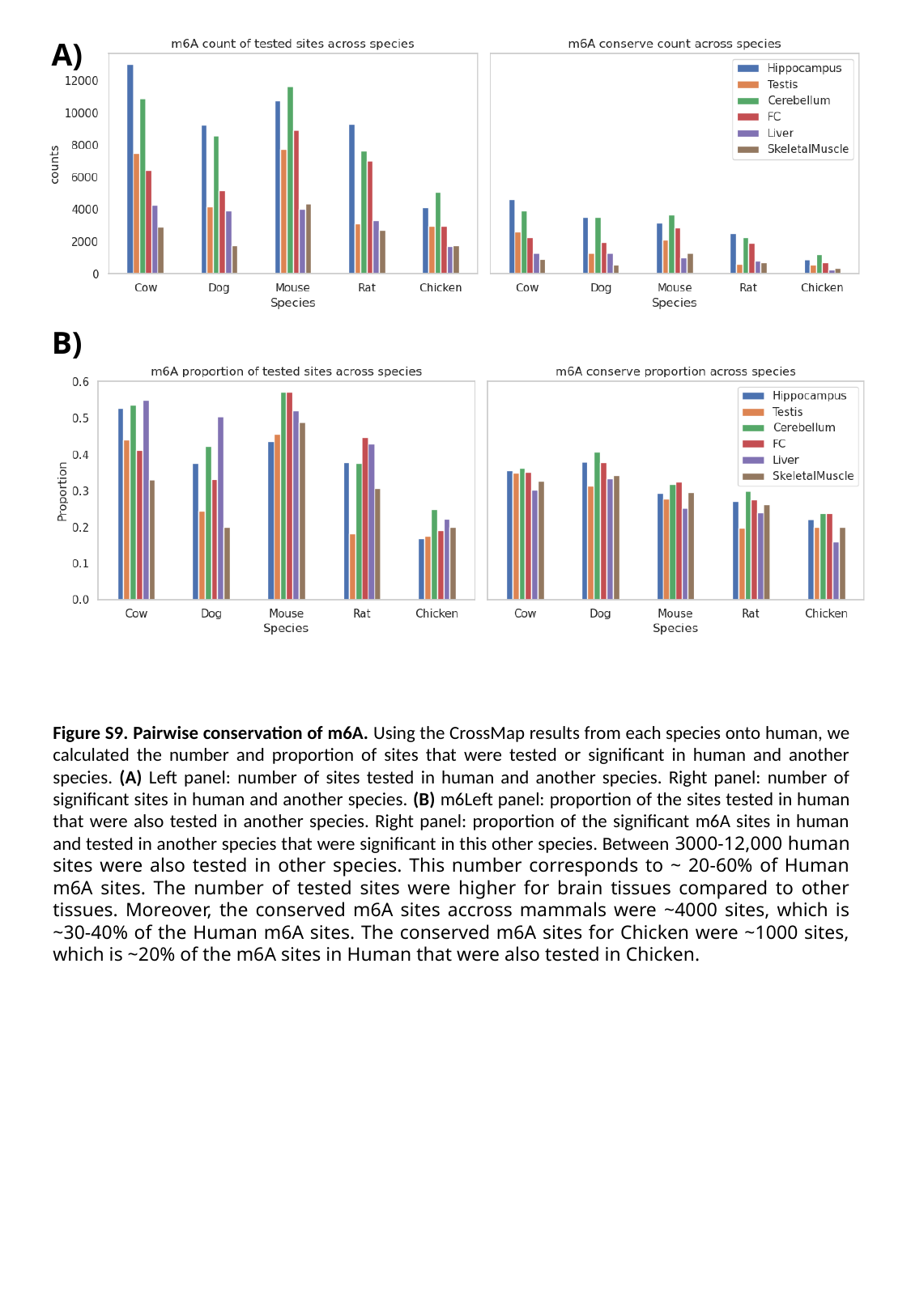

A)
B)
Figure S9. Pairwise conservation of m6A. Using the CrossMap results from each species onto human, we calculated the number and proportion of sites that were tested or significant in human and another species. (A) Left panel: number of sites tested in human and another species. Right panel: number of significant sites in human and another species. (B) m6Left panel: proportion of the sites tested in human that were also tested in another species. Right panel: proportion of the significant m6A sites in human and tested in another species that were significant in this other species. Between 3000-12,000 human sites were also tested in other species. This number corresponds to ~ 20-60% of Human m6A sites. The number of tested sites were higher for brain tissues compared to other tissues. Moreover, the conserved m6A sites accross mammals were ~4000 sites, which is ~30-40% of the Human m6A sites. The conserved m6A sites for Chicken were ~1000 sites, which is ~20% of the m6A sites in Human that were also tested in Chicken.

### Slide 10
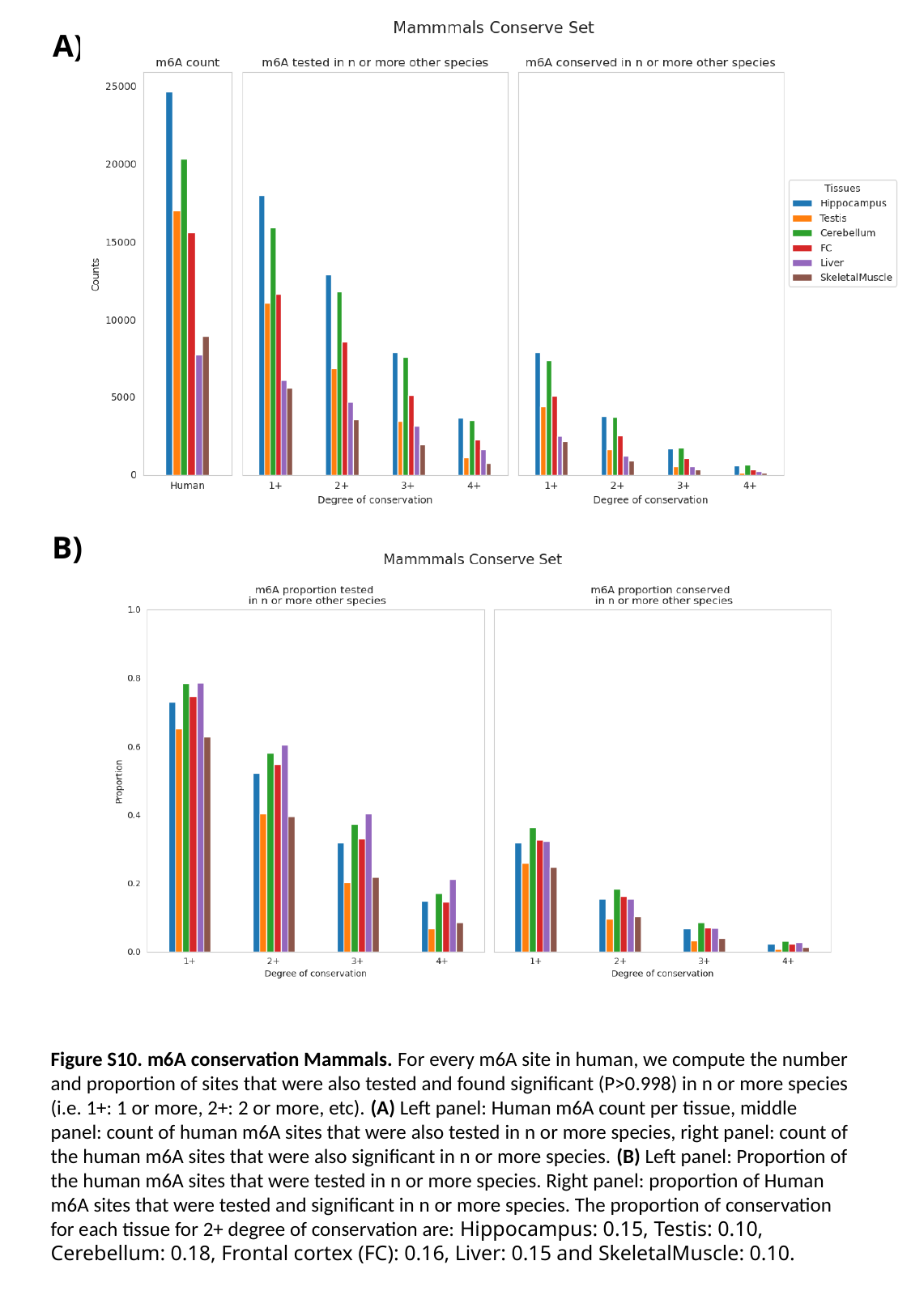

A)
B)
Figure S10. m6A conservation Mammals. For every m6A site in human, we compute the number and proportion of sites that were also tested and found significant (P>0.998) in n or more species (i.e. 1+: 1 or more, 2+: 2 or more, etc). (A) Left panel: Human m6A count per tissue, middle panel: count of human m6A sites that were also tested in n or more species, right panel: count of the human m6A sites that were also significant in n or more species. (B) Left panel: Proportion of the human m6A sites that were tested in n or more species. Right panel: proportion of Human m6A sites that were tested and significant in n or more species. The proportion of conservation for each tissue for 2+ degree of conservation are: ​Hippocampus: 0.15, Testis: 0.10, Cerebellum: 0.18, Frontal cortex (FC): 0.16, Liver: 0.15 and SkeletalMuscle: 0.10.

### Slide 11
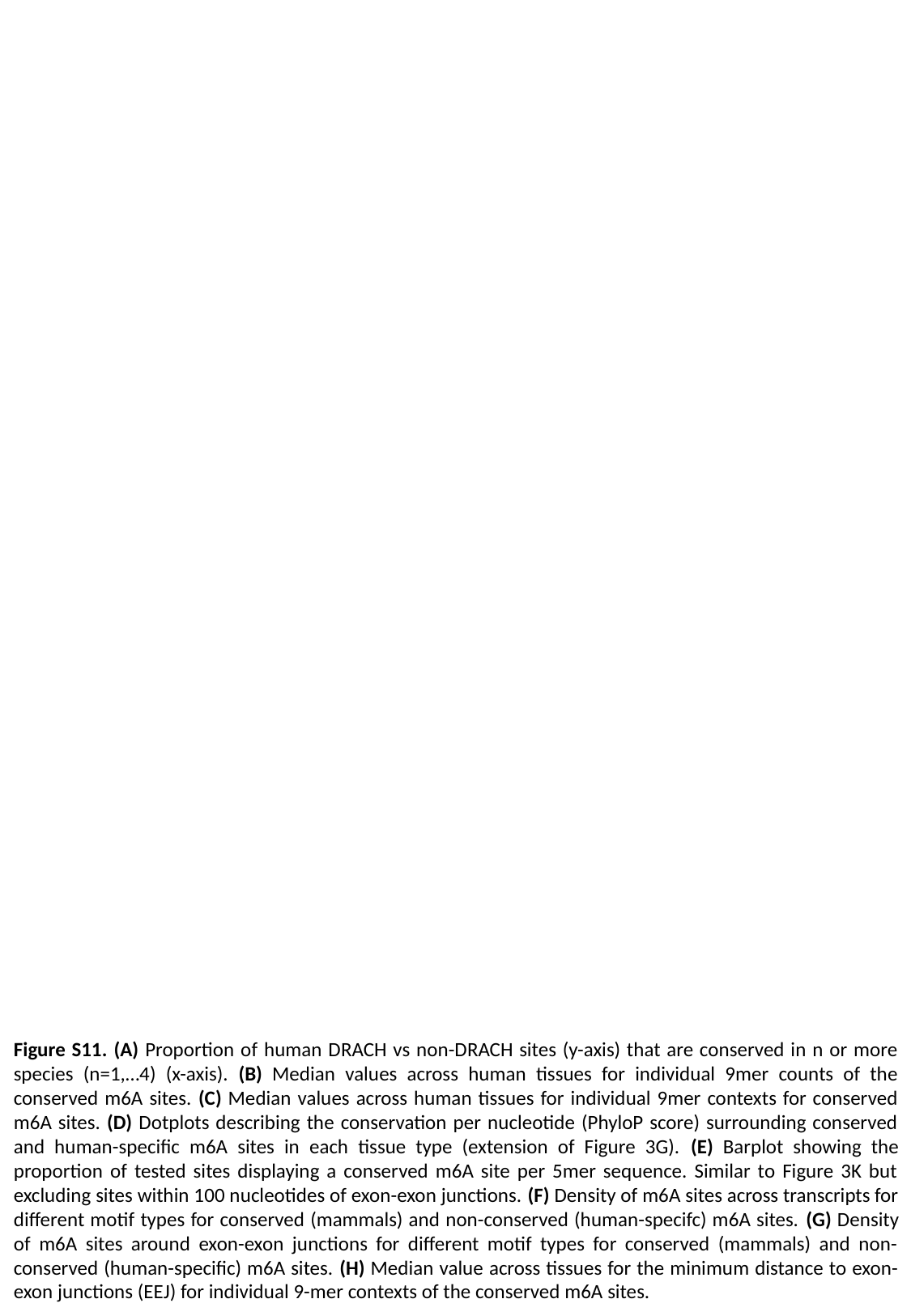

Figure S11. (A) Proportion of human DRACH vs non-DRACH sites (y-axis) that are conserved in n or more species (n=1,…4) (x-axis). (B) Median values across human tissues for individual 9mer counts of the conserved m6A sites. (C) Median values across human tissues for individual 9mer contexts for conserved m6A sites. (D) Dotplots describing the conservation per nucleotide (PhyloP score) surrounding conserved and human-specific m6A sites in each tissue type (extension of Figure 3G). (E) Barplot showing the proportion of tested sites displaying a conserved m6A site per 5mer sequence. Similar to Figure 3K but excluding sites within 100 nucleotides of exon-exon junctions. (F) Density of m6A sites across transcripts for different motif types for conserved (mammals) and non-conserved (human-specifc) m6A sites. (G) Density of m6A sites around exon-exon junctions for different motif types for conserved (mammals) and non-conserved (human-specific) m6A sites. (H) Median value across tissues for the minimum distance to exon-exon junctions (EEJ) for individual 9-mer contexts of the conserved m6A sites.
